# Grasping with a twist: Dissociating action goals from motor actions in human frontoparietal circuits

**DOI:** 10.1101/2023.01.02.522486

**Authors:** Guy Rens, Teresa D. Figley, Jason P. Gallivan, Yuqi Liu, Jody C. Culham

## Abstract

In daily life, prehension is typically not the end goal of hand-object interactions but a precursor for manipulation. Nevertheless, functional MRI (fMRI) studies investigating manual manipulation have primarily relied on prehension as the end goal of an action. Here, we used slow event-related fMRI to investigate differences in neural activation patterns between prehension in isolation and prehension for object manipulation. Sixteen participants were instructed either to simply grasp the handle of a rotatable dial (isolated prehension) or to grasp and turn it (prehension for object manipulation). We used representational similarity analysis to investigate whether the experimental conditions could be discriminated from each other based on differences in task-related brain activation patterns. We also used temporal multivoxel pattern analysis to examine the evolution of regional activation patterns over time. Importantly, we were able to differentiate isolated prehension and prehension for manipulation from activation patterns in the early visual cortex, the caudal intraparietal sulcus, and the superior parietal lobule. Our findings indicate that object manipulation extends beyond the putative cortical grasping network (anterior intraparietal sulcus, premotor and motor cortices) to include the superior parietal lobule and early visual cortex.

**Significance statement:** A simple act such as turning an oven dial requires not only that the central nervous system encode the initial state (starting dial orientation) of the object but also the appropriate posture to grasp it in order to achieve the desired end state (final dial orientation) and the motor commands to achieve that state. Using advanced temporal neuroimaging analysis techniques, we reveal how such actions unfold over time and how they differ between object manipulation (turning a dial) vs. grasping alone. We find that a combination of brain areas implicated in visual processing and sensorimotor integration can distinguish between the complex and simple tasks during planning, with neural patterns that approximate those during the actual execution of the action.

## 1. Introduction

The hand is central in physical interactions with our environment. Typical hand-object interactions consist of sequential phases, starting with reaching and ending with object manipulation (Castiello, 2005). Electrophysiological studies in macaques have implicated a frontoparietal network in hand-object interactions (for a review see Gerbella et al., 2017). Previous functional MRI (fMRI) research (for a review see Errante et al., 2021) has identified a similar network in humans. This network comprises a dorsomedial pathway, consisting of the superior parietal occipital cortex (SPOC, corresponding to V6/V6A) and the dorsal premotor cortex (PMd), and a dorsolateral pathway, consisting of anterior intraparietal sulcus (aIPS) and ventral premotor cortex (PMv).

While past human neuroimaging studies revealed the neural substrates of grasping, most treated prehension as the end goal, not as a step towards meaningful hand-object interactions. In contrast, real-world hand-object interactions typically involve grasping only as a prelude to subsequent actions such as manipulating or moving objects. Importantly, previous studies have shown that the final action goal shapes prehension: Initial prehension strategies are affected by the goal in the ‘end-state comfort’ effect (Rosenbaum et al., 1990) and in actions like tool use (Comalli et al., 2016). Moreover, the end goal affects brain responses during action planning and prehension (Fogassi et al., 2005; Gallivan et al., 2016a).

The purpose of the current study was to investigate how isolated and sequential actions unfold differently on a moment-to-moment basis. The temporal unfolding of brain activation during actions has been largely overlooked (or only studied with EEG, without localizing the specific brain regions involved, e.g., Guo et al., 2019). Most studies that have used multivoxel pattern analysis (MVPA) to investigate hand actions have averaged data within time bins for planning and execution (e.g., Gallivan et al., 2011). Notably, several studies have revealed the temporal unfolding of MVPA grasping representations for sequential timepoints (Ariani et al., 2018a; Gallivan et al., 2013). In addition, one study examined isolated (grasping an object) vs. sequential (grasping to move an object to one of two locations) actions, showing that activation patterns could be discriminated across the grasping network even during action planning (Gallivan et al., 2016).

Here we examined (univariate) activation levels and (multivariate) activation patterns (using representational similarity analysis, RSA (Kriegeskorte, 2008). Moreover, we also used a new methodological approach, temporal MVPA (tMVPA; Ramon et al., 2015; Vizioli et al., 2018) to examine the representation similarities across trials for the same and different points in time, separately for isolated vs. sequential actions.

We measured brain activation using fMRI while participants performed a motor task consisting of either simple grasping of a dial (with two possible initial orientations) or grasping followed by rotation of the dial (clockwise or counterclockwise), as one might turn an oven dial (See Figure 1). Based on earlier findings that brain regions within the grasping network plan the full action sequence, and not just the initial grasp (see Gallivan et al., 2016), we expected that during the plan phase, as well as the execute phase, tMVPA would reveal representations of the task (grasp vs. turn). Given that visual orientation (Kamitani & Tong, 2005), surface/object orientation (Rice et al., 2007; Shikata et al., 2001; Valyear et al., 2006) and grip orientation (Monaco et al., 2011) are represented in early visual cortex, the caudal intraparietal sulcus (cIPS), and reach-selective cortex (SPOC), respectively, we predicted that the representation of orientation would be biased to the start orientation early in planning, with a greater emphasis on end orientation as the movement progressed. Our approach with tMVPA allowed us to examine, across different regions, how representations unfolded over time for isolated vs. sequential actions. Specifically, we expected that regions sensitive to the kinematics of executed actions (e.g., M1 and S1) would only show highly similar representations during action execution; whereas, regions involved in more abstract features of action planning (e.g., aIPS for coding object shape) would show similar representations across planning and execution.

**Figure 1.**
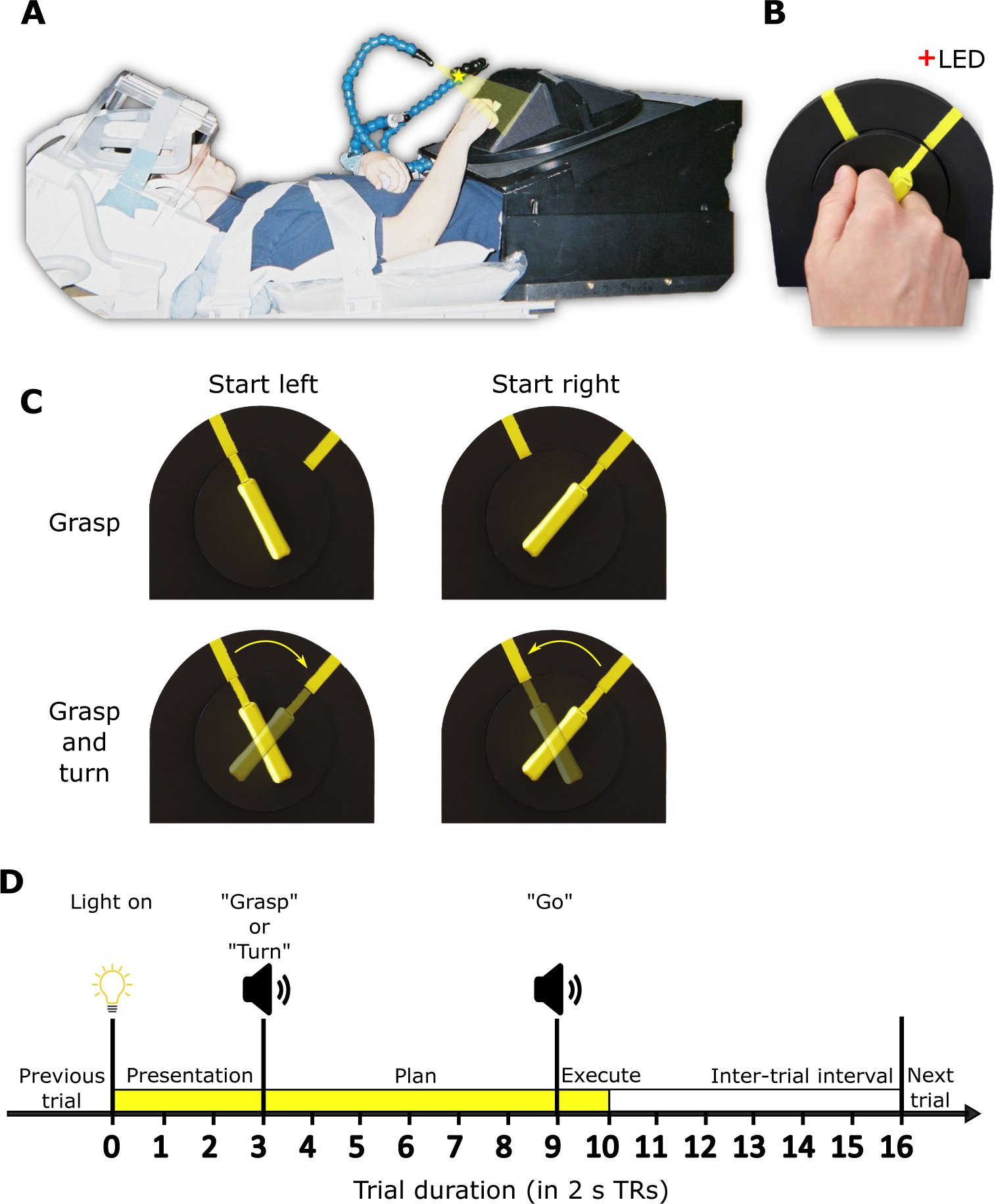
Experimental set-up and design. **1A.** Picture of participant set-up in the fMRI scanner shown from side view. **1B.** Close up of participant using the instructed grip to grasp the experimental stimulus, i.e., rotatable dial with a handle. **1C.** Experimental conditions in a two (start orientation: left or right) by two (action: grasp or turn) design. **2D.** Timing of each event-related trial. Trials began with the stimulus being illuminated (TR 0-3) indicating the start of the preparation phase. At TR 3 participants received task instructions indicating the start of the planning phase (TR 3-9). At TR 9 participants received the “go” cue indicating onset of action execution (TR 9-10). After 2 s (1 TR) the light was turned off and the intertrial interval was initiated (TR 10-16)

## 2. Methods

### 2.1 Participants

Data from sixteen right-handed volunteers was utilized in the analysis (7 males, 9 females, mean age: 24.4 years). Participants were recruited from Western University (London, Ontario, Canada) and provided informed consent in accordance with procedures approved by the University’s Health Sciences Research Ethics Board. Data from an additional two subjects (one male and one female) was collected but excluded due to excessive motion artifacts (see 2.6).

### 2.2 Setup and apparatus

The experimental setup is illustrated in Figure 1A-B. Participants lay supine in a 3-Tesla MRI scanner with the head and head coil tilted approximately 30° to allow for direct viewing without mirrors of a manipulandum positioned above the participant’s hips. The manipulandum consisted of a black rotatable dial (9-cm diameter; Figure 1B) with a yellow rectangular handle (5-cm length x 1-cm width x 2-cm depth). The dial was mounted on a black surface. The black surface was positioned such that the dial was approximately perpendicular to the subject’s line of gaze and comfortably within reach of their right arm. Two yellow markers were put on the black surface (Figure 1B) indicating the start and end positions for turning the dial. A grey platform (not shown in Figure 1A) was positioned above the participant’s lower torso serving as the home/resting position for the right arm between trials. Participants’ upper arms were braced above their torsos and just above their elbows (Figure 1A) to limit movement of the shoulder, which can induce motion artifacts in fMRI signals. As such, participants could only rely on elbow flexion/extension and forearm rotation to perform the experimental task. Considering these constraints, the position and orientation of the dial were adjusted for each participant to optimize participant comfort during task performance and ensure the dial remained fully visible. The position of the yellow markers on the black surface was adjusted for each participant individually so that dial rotation would not exceed 80% of the participant’s maximum range of motion when turning the dial clockwise or counterclockwise.

During the experiment, the dial was illuminated from the front by a bright yellow Light Emitting Diode (LED) attached to flexible plastic stalks (Figure 1A; Loc-Line, Lockwood Products, Lake Oswego, OR). A dim red LED (masked by a 0.1° aperture) was positioned approximately 15° of visual angle above the rotatable dial and just behind it to provide a fixation point for participants (Figure 1A-B). Experimental timing (see below) and lighting were controlled with in-house MATLAB scripts (The MathWorks Inc., Natick, MA).

### 2.3 Experiment design and timing

#### Behavioral task

This experiment was a 2 (starting orientation: left or right) x 2 (action: grasp or turn) delayed movement paradigm (Figure 1C). For each trial, the dial would appear in one of the two yellow-marked starting positions. The grasp condition consisted of reaching towards the dial and squeezing it between the middle phalanges of the index and middle fingers (trial end position shown in Figure 1A; hand shape during grasp shown in Figure 1B). After grasp completion, participants returned their arm back to the home position. In the turn condition, participants performed the same reach-to-grasp action but would then subsequently rotate the dial clockwise or counterclockwise after they grasped the dial in the left or right start position, respectively (Figure 1C). We decided on the index-middle finger grip instead of a more natural variant of precision grasp as this would ensure the grip would be highly similar in all conditions and limit changes to grip angle to optimize end-state comfort (e.g., putting the right-hand thumb further up when planning to turn clockwise and further down when planning to turn the dial counterclockwise). This avoids contaminating neural activation with low-level sensorimotor confounds such as digit positioning. Participants were instructed to keep the timing of all movements as similar as possible, such that the right hand reached from and returned to the home position at the same time (see next paragraph ‘Trial Design’). To isolate the visuomotor planning response from the visual and motor execution responses, we used a slow event-related paradigm with 32-s trials, consisting of three phases: ‘Presentation’, ‘Plan’ and ‘Execute’ (see Figure 1D). We adapted this paradigm from previous fMRI studies that successfully isolated delay-period activity from transient neural responses following the onset of visual stimuli and movement execution (Beurze et al., 2007, 2009; Pertzov et al., 2011). Furthermore, using this paradigm in previous work from our group, we were able to successfully isolate and decode planning-related neural activation prior to action execution (Gallivan et al., 2011; Gallivan, McLean, et al., 2013).

#### Trial design

Before each trial, subjects were in complete darkness except for the fixation LED upon which participants were instructed to maintain their gaze. The trial began with a 6-s (3 TRs of 2 s each) Presentation phase in which the illumination LED lit up the rotatable dial. After the Presentation phase, the 12-s (6-TR) Plan phase was initiated with a voice cue (0.5-s duration) saying either ‘Grasp’ or ‘Turn’ to instruct the upcoming action to the participant. Participants could see the object during the presentation phase, thus perceiving the starting orientation of the yellow handle. However, they were instructed to only begin the action after they received the ‘Go’ cue. After the Plan phase, a 2-s (1-TR) Execute phase began with a 0.5-s beep (the ‘Go’ cue’) cueing participants to initiate and execute the instructed action. Performing the entire action, consisting of reaching and grasping (with or without dial turning), took approximately 2 s. The dial remained illuminated for 2 s after the ‘Go’ cue allowing visual feedback during action execution. After the 2-s Execution phase, the illumination LED was turned off, cueing participants to let go of the dial and return their hand back to the home position. Participants remained in this position for 12 s (6 TRs) in the dark (i.e., the intertrial interval) to allow the BOLD response to return back to baseline before the next trial would be initiated.

#### Functional runs

Within each functional run, each of the four trial types (grasp: left or right – turn: from left to right or right to left) was presented five times in a pseudo-randomized manner for a total of 20 trials. Each participant performed eight functional runs, yielding a total of 40 trials/condition (160 trials in total). Each functional run took between 10 and 11 minutes. For each participant, trial orders were counterbalanced across all functional runs so that each trial type was preceded and followed equally often by every trial type (including the same trial type) across the entire experiment. During testing, the experimenter was positioned next to the set-up to ensure that the dial was in the correct position before each trial, and manually adjust it if needed. In addition, the experimenter could visually check whether the participant performed the trial correctly. Incorrectly trials were defined as any trial where the participant did not perform the task as instructed: for instance, turning instead of grasping, using incorrect grips, failing to turn in a smooth manner or having the dial slip from the fingers.

Participants performed a separate practice session before the actual experiment to familiarize them with the motor task and ensure proper execution. The experimental session took approximately three hours and consisted of preparation (i.e., informed consent, MRI safety, placing the participant in the scanner and setting up the behavioral task), eight functional runs and one anatomical scan. The anatomical scan was collected between the fourth and fifth functional runs to give participants a break from the task.

### 2.4 MRI acquisition

Imaging was performed using a 3-Tesla Siemens TIM MAGNETOM Trio MRI scanner at the Robarts Research Institute (London, ON, Canada). The T1-weighted anatomical image was collected using an ADNI MPRAGE sequence (time to repetition (TR) = 2300 ms, time to echo (TE) = 2.98 ms, field of view = 192 mm x 240 mm x 256 mm, matrix size = 192 x 240 x 256, flip angle = 9°, 1-mm isotropic voxels). Functional MRI volumes sensitive to the blood oxygenation level-dependent (BOLD) signal were collected using a T2*-weighted single-shot gradient-echo echo-planar imaging (EPI) acquisition sequence (TR = 2000 ms, slice thickness = 3 mm, in-plane resolution = 3 mm x 3 mm, TE = 30 ms, field of view = 240 mm x 240 mm, matrix size = 80 x 80, flip angle = 90°, and acceleration factor (integrated parallel acquisition technologies, iPAT) = 2 with generalized auto-calibrating partially parallel acquisitions (GRAPPA) reconstruction. Each volume comprised 34 contiguous (i.e., with no gap) oblique slices acquired at an approximate 30° caudal tilt with respect to the anterior-to-posterior commissure (ACPC) plane, providing near whole brain coverage. We used a combination of parallel imaging coils to achieve a good signal to noise ratio and to enable direct viewing of the rotatable dial without mirrors or occlusion. Specifically, we placed the posterior half of the 12-channel receive-only head coil (6-channels) beneath the head and tilted it at an angle of approximately 20°. To increase the head tilt to approximately 30°, we put additional foam padding below the head. We then suspended a 4-channel receive-only flex coil over the forehead (Figure 1A).

### 2.5 fMRI anatomical data processing

All fMRI preprocessing was performed in BrainVoyager version 22 (Brain Innovation, Maastricht Netherlands). For the present study, we defined regions of interest (ROIs) in surface space instead of volumetric space as it has been shown that cortical alignment improves group results by reducing individual differences in sulcal locations (Fischl et al., 1999; Frost & Goebel, 2012). As such, we performed surface reconstruction (mesh generation) and cortex-based alignment. Given that we relied on the recommended approach and standard settings of BrainVoyager 22, these steps are explained only in brief.

#### Folded mesh generation for each participant

In volumetric space we performed the following steps. First, we corrected for intensity inhomogeneity. Then, we rotated the anatomical data in ACPC space, due to the experimental head tilt, and normalized to Montreal Neurological Institute (MNI) space. Next, we excluded the subcortical structures (labelled as white matter) and the cerebellum (removed from anatomical) from mesh generation. We then defined the boundaries between white matter and gray matter and between gray matter and cerebrospinal fluid. Finally, we created a folded mesh (surface representation) of only the left hemisphere after removing topologically incorrect bridges (Kriegeskorte & Goebel, 2001). We decided to only investigate the left hemisphere because the motor task involved the right hand only, which previous work has shown predominantly activates the left (contralateral) hemisphere (Cavina-Pratesi et al., 2010, 2018; Gallivan et al., 2011; Gallivan, McLean, et al., 2013).

#### Standardized folded mesh generation for each participant

Briefly, folded meshes created from anatomical files often result in different numbers of vertices between participants. To facilitate cortex-based alignment between participants, folded meshes were first transformed into high-resolution standardized folded meshes. Each folded mesh was first morphed into a spherical representation by smoothing (thus removing differences between sulci and gyri) and correcting for distortion. The spherical representation of each participant mesh was then mapped to a high-resolution standard sphere to create a high-resolution standardized spherical representation of the participant mesh. The vertex position information of the original participant folded (not spherical) mesh was then used to generate a standardized folded mesh for each participant.

#### Cortex-based alignment

Cortex-based alignment was performed following the approach of Frost & Goebel (2012) and Goebel et al. (2006): We aligned all individual standard meshes to a dynamically generated group average target mesh. Before aligning to the dynamic group average, we performed pre-alignment (i.e., rigid sphere alignment). The actual steps of cortex-based alignment generate a dynamic group average (a surface mesh based on all individual meshes) and sphere-to-sphere mapping files for each participant that enable transporting the functional data from each individual to the dynamic group average. Inverse sphere-to-sphere mapping files were also generated, which allows transporting of data (such as regions of interest) from the dynamic group average back to individual meshes.

### 2.6 fMRI functional data processing

#### General preprocessing

All functional runs were screened for motion and magnet artifacts by examining the movement time courses and motion plots created with the motion correction algorithms (three translation and three rotation parameters). Data from a run was discarded if translation exceeded 1 mm or rotation exceeded 1° between successive volumes. Based on this screening, all data from two participants and one run from an additional participant was discarded. Examination of the remaining data indicated negligible motion artifacts in time courses (as will be shown in later figures).

Functional runs were co-registered with the anatomical data using boundary-based registration (Greve & Fischl, 2009). We subsequently performed slice-scan time correction, motion correction and MNI normalization. Next, we performed linear-trend removal and temporal high-pass filtering (using a cut-off of three sine and cosine cycles on the fast Fourier transform of the time courses).

#### Volumetric to surface-based time courses

Volumetric time courses were first aligned with the respective participant’s anatomical scan using boundary-based registration to ensure optimal alignment (Greve & Fischl, 2009). Volumetric time courses were then transformed to the respective participant’s standardized folded mesh using depth integration along the vertex normal (sampling from -1 to 3 mm relative to the gray-white matter boundary). Once functional runs were transformed into surface space, they were spatially smoothed. Finally, functional runs in individual standard mesh space, could then be mapped onto the dynamic group average mesh using sphere-to-sphere mapping files that were generated during cortex-based alignment. This approach enabled us to model the averaged brain activation across all participants onto the group mesh and define ROIs based on hotspots of strongest activations in surface space.

### 2.7 Region of interest-based analysis

For the current project, we performed a ROI-based analysis of cortical regions within the hemisphere contralateral to the acting hand at the peak foci for task-related activity. To allow for better intersubject consistency and statistical power, we utilized cortex-based alignment for defining ROIs (Frost & Goebel, 2012; Goebel et al., 2006). We were primarily interested in cortical areas that constitute the cortical grasping and reaching networks that have been investigated in previous studies of our group (Cavina-Pratesi et al., 2018) and others (Ariani et al., 2018). As in previous projects (e.g., Gallivan et al., 2018; Fabbri et al., 2016), we considered only the contralateral hemisphere as it is well-established that hand-object actions are strongly lateralized and elicit only limited activation in the ipsilateral hemisphere (Culham et al., 2006). Moreover, as our analysis of the left cerebral hemisphere already included 17 regions and we performed multiple analyses, some at many time points, we wanted to limit the complexity of the results and find a balance between statistical rigour and statistical power.

We chose an ROI approach rather than a whole-brain cortex-based searchlight to optimize statistical power and computational processing time (Etzel et al., 2013; Frost & Goebel, 2012; Kriegeskorte, 2008). Due to the large number of comparisons performed in a searchlight analysis, only the most robust effects survive the correction for multiple comparisons (Kriegeskorte, 2008; Kriegeskorte & Kievit, 2013). Moreover, tMVPA relies on single-trial correlations which are computationally very intensive. Performing this analysis on the voxel-based searchlight level would likely take multiple months.

### 2.8 Defining regions of interest

To localize our regions of interest on the group mesh, we applied a general linear model (GLM) on our data in surface space. Predictors of interest were created from boxcar functions convolved with the two-gamma haemodynamic response function (HRF). We aligned a boxcar function with the onset of each phase of each trial within each run of each participant, with a height dependent upon the duration of each phase. We used 3 TRs for the presentation phase, 6 TRs for the Plan phase and 1 TR for the Execution phase. The six motion parameters (three translations and three rotations) were added as predictors of no interest. Each incorrect trial was also assigned a unique predictor of no interest. All regression coefficients (beta weights) were defined relative to the baseline activity during the intertrial interval. In addition, time courses were converted to percent signal change before applying the random effects GLM (RFX-GLM).

To specify ROIs for our analyses, we searched for brain areas, on the group level, involved in the experimental task. We contrasted brain activation for all three phases with baseline. This contrast was performed across all conditions to ensure that ROI selection based on local activation patterns was not biased by differences between specific conditions (e.g., grasp versus turn). Specifically, the contrast was: {Presentation [Grasp _left_ + Grasp _right_ + Turn _left to right_ + Turn _right to left_] + Plan [Grasp _left_ + Grasp _right_ + Turn _left to right_ + Turn _right to left_] + Execute [Grasp _left_ + Grasp _right_ + Turn _left to right_ + Turn _right to left_]} > baseline. This contrast enabled us to identify regions that showed visual and/or motor activation associated with the task.

In BrainVoyager, area selection on a surface generates a hexagon surrounding the selected vertex. We decided on an area selection size of 100 (arbitrary units) as this value provided a good balance between inclusion of a sufficient number of vertices/voxels around each hotspot for multivariate analyses and avoiding overlap between ROIs (especially in the parietal lobe).

### 2.9 Regions of interest

Selected regions of interest (Figure 2) were defined in the left hemisphere. Most regions of interest were defined using the contrast above (all conditions > baseline), with two exceptions, as detailed below.

**Figure 2.**
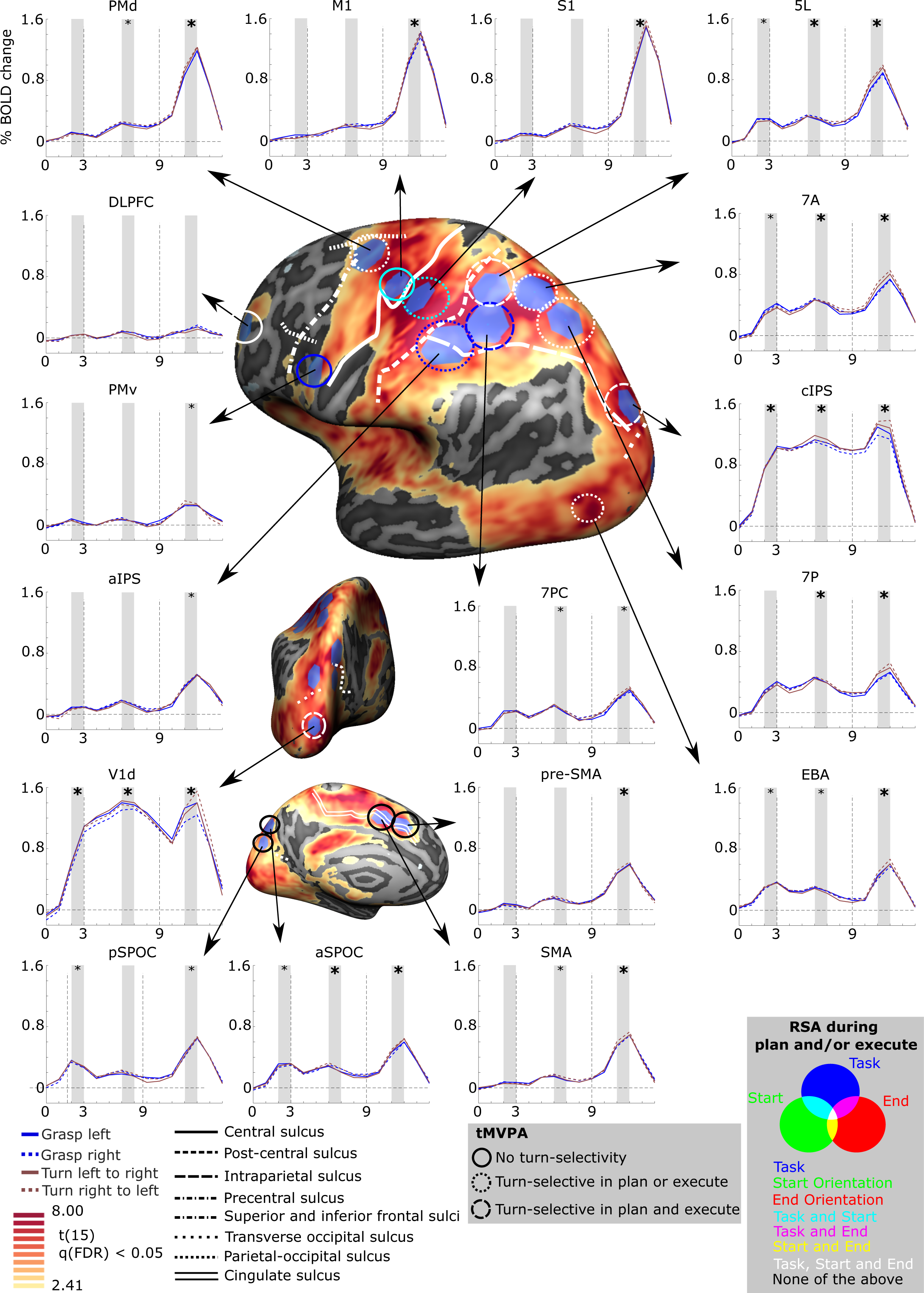
Brain areas selected for multivariate analysis based on a univariate contrast. Cortical areas that exhibited larger responses during the experimental trials [(presentation + plan + execute) > baseline] are shown in yellow to red activation. Results calculated across all participants (RFX GLM) are displayed on the dynamic group average surface across participants. The selected ROIs were selected in BrainVoyager software, which uses a hexagonal selection area, and then transformed to individual volumetric MNI space. Each ROI is linked to the group average of corresponding % signal change of BOLD activity (Y-axis) over time (X-axis; each X-ticks represents one TR of 2 s, based on a deconvolution analysis) for each of the four conditions. Vertical dashed lines on the graphs indicate start of the planning phase (TR 3) and the execution phase (TR 9). For the time courses, statistical analysis was done for the shaded areas (e.g., average timepoint 2 and 3) averaged across the 4 conditions. If significantly > 0, when Bonferroni corrected within ROI-only, a small asterisk is depicted. If significantly > 0, when Bonferroni corrected for all comparisons, a large asterisk is depicted. On the surface brain, sulcal landmarks are denoted by white lines (stylized according to the corresponding legend). ROI acronyms are spelled out in the methods. tMVPA results are indicated using different types of circles (e.g., dashed vs. solid). RSA results are shown by color-coding the tMVPA circles using a Venn diagram. Note that the RSA and tMVPA results are discussed in the results section as well as in Figure 5 and Figure 6.

First, the involvement of the cortical grasping network (Gerbella et al., 2017) was investigated by including sensorimotor and visuomotor regions (Ariani et al., 2018; Cavina-Pratesi et al., 2018; Fabbri et al., 2016; Gallivan et al., 2011; Gallivan, McLean, et al., 2013).

- Primary motor cortex (M1): Hotspot of strongest activation near the ‘hand knob’ landmark (Gallivan et al., 2013; Yousry et al., 1997).
- Dorsal premotor cortex (PMd): Hotspot of strongest activation near the junction of the precentral sulcus and the superior frontal sulcus (Picard & Strick, 2001; Pilacinski et al., 2018).
- Ventral premotor cortex (PMv): Hotspot of strongest activation inferior and posterior to the junction of the inferior frontal sulcus) (Tomassini et al., 2007).
- Primary somatosensory cortex (S1): Hotspot of strongest activation anterior to the anterior intraparietal sulcus, encompassing the postcentral gyrus and sulcus (Gallivan et al., 2011; Gallivan, McLean, et al., 2013)
- Anterior intraparietal sulcus (aIPS): Hotspot of strongest activation directly at the junction of the intraparietal sulcus and the postcentral sulcus (Culham et al., 2003).
- Caudal intraparietal sulcus (cIPS): Hotspot of strongest activation on the lateral side of the brain, anterior and superior to the junction between the intraparietal sulcus and the posterior occipital sulcus (Beurze et al., 2009; Grefkes & Fink, 2005). Comparisons with the Julich atlas (Richter et al., 2019) suggested overlap with hIP7 and/or hP01.
- Anterior superior parietal occipital sulcus (aSPOC): Hotspot of strongest activation on the medial side of the brain, anterior and superior to the parietal-occipital sulcus (Cavina-Pratesi et al., 2010), thought to correspond to area V6A(Pitzalis et al., 2015).
- Posterior superior parietal occipital sulcus (pSPOC): Hotspot of strongest activation on the medial side of the brain, posterior and inferior to the parietal-occipital sulcus (Cavina-Pratesi et al., 2010). thought to correspond to area V6 (Pitzalis et al., 2015).

Second, the following medial and frontal regions were selected due to their involvement in motor planning and decision making (Ariani et al., 2015; Badre & Nee, 2018; Cavina-Pratesi et al., 2018).

- Dorsolateral prefrontal cortex (DLPFC): Hotspot of strongest activation near the middle frontal gyrus (Mylius et al., 2013).
- Supplementary motor area (SMA): hotspot of strongest activation adjacent to the medial end of the cingulate sulcus and posterior to the plane of the anterior commissure (Picard & Strick, 2001).
- Pre-supplementary motor area (pre-SMA): Hotspot of strongest activation superior to the cingulate sulcus, anterior to the plane of the anterior commissure and anterior and inferior to the hotspot of strongest activation selected for SMA (Picard & Strick, 2001).

Third, we included the superior parietal lobule (SPL) due to the extensive activation evoked by our task and in hand actions more generally (Ariani et al., 2018a; Cavina-Pratesi et al., 2018). We defined the SPL as the area on the lateral/superior side of the brain that is bordered anteriorly by the postcentral sulcus, inferiorly by the intraparietal sulcus and posteriorly by the parietal-occipital sulcus (Scheperjans, et al., 2008a; Scheperjans et al., 2008b). Due to the large swathe of activation evoked by our contrast, we defined four ROIs within the SPL based on their relative position to each other (to ensure minimal overlap) and the anatomical landmarks bordering the SPL. We decided on four ROIs as well as their names based on Scheperjans et al. (2008a).

- 7PC: Hotspot of strongest activation located on the posterior wall of the postcentral sulcus and superior to the intraparietal sulcus. Given we defined post-aIPS as well, 7PC was also defined as superior to post-AIPS.
- 5L: Hotspot of strongest activation located just posterior to the postcentral sulcus and superior to area 7PC.
- 7P: Hotspot of strongest activation superior to the intraparietal sulcus and anterior to the parietal-occipital sulcus.
- 7A: Hotspot of strongest activation in the postcentral gyrus, superior to 7PC, posterior to 5L and anterior to 7P.

Finally, two visual regions were selected from a probabilistic functional atlas that utilized cortex-based alignment (Rosenke et al., 2021), to which we aligned our participant surface meshes.

- Dorsal primary visual cortex (V1d): Given that primary visual cortex (V1) responds to visual orientation (Kamitani & Tong, 2005b) we wanted to investigate its response here. Because our target objects (and participants hands) fell within the lower visual field (below the fixation point), they would stimulate the dorsal divisions of early visual areas (Wandell et al., 2007). As such, we investigated only the dorsal division of V1, V1d, using the Rosenke atlas.
- Extrastriate body area (EBA): We were also interested in examining the response of the extrastriate body area, which has been implicated not only in the visual perception of bodies (Greenfield et al., 1996) but also in computing goals during action planning (Astafiev et al., 2004; Zimmermann et al., 2016). Because the EBA is not easy to distinguish from nearby regions of the lateral occipitotemporal cortex, we utilized the EBA from the Rosenke atlas. Briefly, this process is nearly identical for the other ROIs. The only difference was that we aligned all individual standard meshes to the template of Rosenke (instead of the dynamic group average). This provided sphere-to-sphere and inverse sphere-to-sphere mapping files. The latter could then be used to transport EBA from the Rosenke template to the individual standard meshes and perform all other steps identically as for the other ROIs. We defined EBA as the combination of all body-selective patches, being OTS-bodies, MTG-bodies, LOS-bodies and ITG-bodies (for the full explanation of these regions see Rosenke et al., 2021)

After ROIs were defined in group surface space, they were transformed to individual surface space, using the inverse transformation files generated during cortex-based alignment and then to individual volumetric MNI space, using depth expansion (inverse of depth integration; see ‘fMRI functional data processing: Volumetric to surface-based time courses’) along the vertex normals (-1 to 3 mm). This approach allowed us to define ROIs on the group surface but extract functional data from the individual volumetric level.

### 2.10 Analysis of functional data

Functional data were extracted to perform deconvolution general linear models (deconvolution GLMs; Hinrichs et al., 2000), representational similarity analysis (Kriegeskorte, 2008) and temporal multivoxel pattern analysis (tMVPA; Vizioli et al., 2018). All analyses described below, excluding initial processing of fMRI data and the univariate analyses (which where both done in BrainVoyager), were done with in-house MATLAB scripts.

#### 2.10.1 Univariate analysis

We used a random-effects GLM with deconvolution to extract the time courses of activation in each ROI during the experimental task. For each experimental condition, we used 15 matchstick predictors (timepoint 0 to 14), the first predictor (“predictor 1”) was aligned with time point zero (Figure 1D; “light on”) and the last predictor (“predictor 15”) was aligned with the drop in signal at the end of the trial (“timepoint 14). Due to this clear drop in signal at timepoint 14, we did not include a predictor (non-existing “predictor 16”) for the last timepoint (existing timepoint 15; see Figure 1). For each functional voxel in each ROI, baseline *z*-normalized estimates of the deconvolved BOLD response, representing the mean-centered signal for each voxel and condition relative to the standard deviation of signal fluctuations, were extracted for all conditions. Voxel activation was then averaged across voxels within each ROI. This provided us with an estimate of the averaged time course of brain activation for each condition for each ROI and for each participant.

Given previous work from our group demonstrating the involvement of our selected ROIs in hand-object interactions (e.g., Fabbri et al., 2016; Gallivan et al., 2009; Monaco et al., 2011), we included univariate time courses for comparison as they indicate changes in the signal and reveal the degree to which differences in activation levels are present (or not). The univariate time courses also demonstrate that our quality-assurance procedures, particularly motion screening, succeeded in generating clean data.

Time courses were generated for each ROI (as shown in Figure 2) separately for each of the four conditions. However, the differences in univariate signals between conditions were small or negligible. Thus, to simply evaluate whether or not there was significant activation during each of the three phases of the trial, we performed statistical comparisons of the average activation levels (collapsed across the four conditions) compared to the intertrial baseline (time 0).

First, for each ROI in each participant we averaged the data across conditions. Second, to reduce the number of statistical comparisons while targeting peak responses, we defined three distinct phases by averaging two datapoints for each. These phases were response (i) to presentation, by averaging timepoint 2 and 3, (ii) to planning, by averaging timepoint 6 and 7, (iii) and to execution by averaging timepoint 11 and 12. Third, we investigated significant changes compared to baseline (i.e., > 0) using one-sample *t* tests. Accordingly, per ROI we performed three one-sample *t* tests. Fourth, we corrected for multiple comparisons. It is well-known that balancing type I and type II errors in fMRI research can be challenging due to the amount of data and number of comparisons involved. Although data between ROIs is independent, the number of comparisons in total across ROIs increase the likelihood of Type I errors. Because of this predicament, we decided to correct the univariate analysis for multiple comparisons using the Bonferroni method in two manners being (a) only including comparisons within ROIs (i.e., 3 comparisons within each ROI thus *alpha* = 0.05/3) and (b) including all comparisons (i.e., 3 comparisons within each of the 17 ROIs thus *alpha* = 0.05/51). The results can be found in Figure 2: a small asterisk indicates that the values are significant against the standard correction (3 comparisons), a larger bold asterisk indicates that the values are significant against the strict correction (51 comparisons). The values in the results reflect mean ± SEM. *P* values will be referred to as *P* and *P*(strict) for the 3-comparison and 51-comparison corrections, respectively. Note that we provide the *p* values corrected for the comparisons. For instance, if we found a p-value of 0.0035, it would be reported as P = 0.0105 (corrected for 3 comparisons) and P(strict) = 0.1785 (corrected for 51 comparisons).

#### 2.10.2 Temporal multivoxel pattern analysis

For tMVPA we relied on the approach developed in Ramon et al. (2015) and Vizioli et al. (2018). Briefly, tMVPA has been developed to investigate the temporal development of neural representations by relying on multivariate analyses with a trial wise approach. The tMVPA methods are schematized in Figure 3. For each voxel of each ROI of each participant, we performed a deconvolution GLM for every trial separately, which was then transformed into BOLD percent signal change by dividing the raw BOLD time course by its mean. Note that for tMVPA we performed deconvolution GLMs for each trial (of the same condition) separately whereas for the univariate analysis and RSA (see below) we ran deconvolution GLMs for each condition (i.e., across same-condition trials). After computing the single-trial activation patterns, we computed single-trial representational dissimilarity matrices (stRDMs; dissimilarity = 1 - *r*) using Pearson correlations for each condition of each ROI of each participant. As shown in Figure 3A, a stRDM is generated by computing dissimilarity between the activation patterns of voxels at each timepoint of a given trial (‘trial m’) with the activation pattern of the same voxels for each of the timepoints for another trial (‘trial n’). This process is iteratively repeated until all unique within-condition trial pairings are run. stRDMs were then averaged across the main diagonal to yield a diagonally symmetric matrix. That is, the data matrix was averaged position-wise with its transpose matrix. We performed this averaging as we were only interested in between-condition differences and not in differences between same-condition trials.

**Figure 3.**
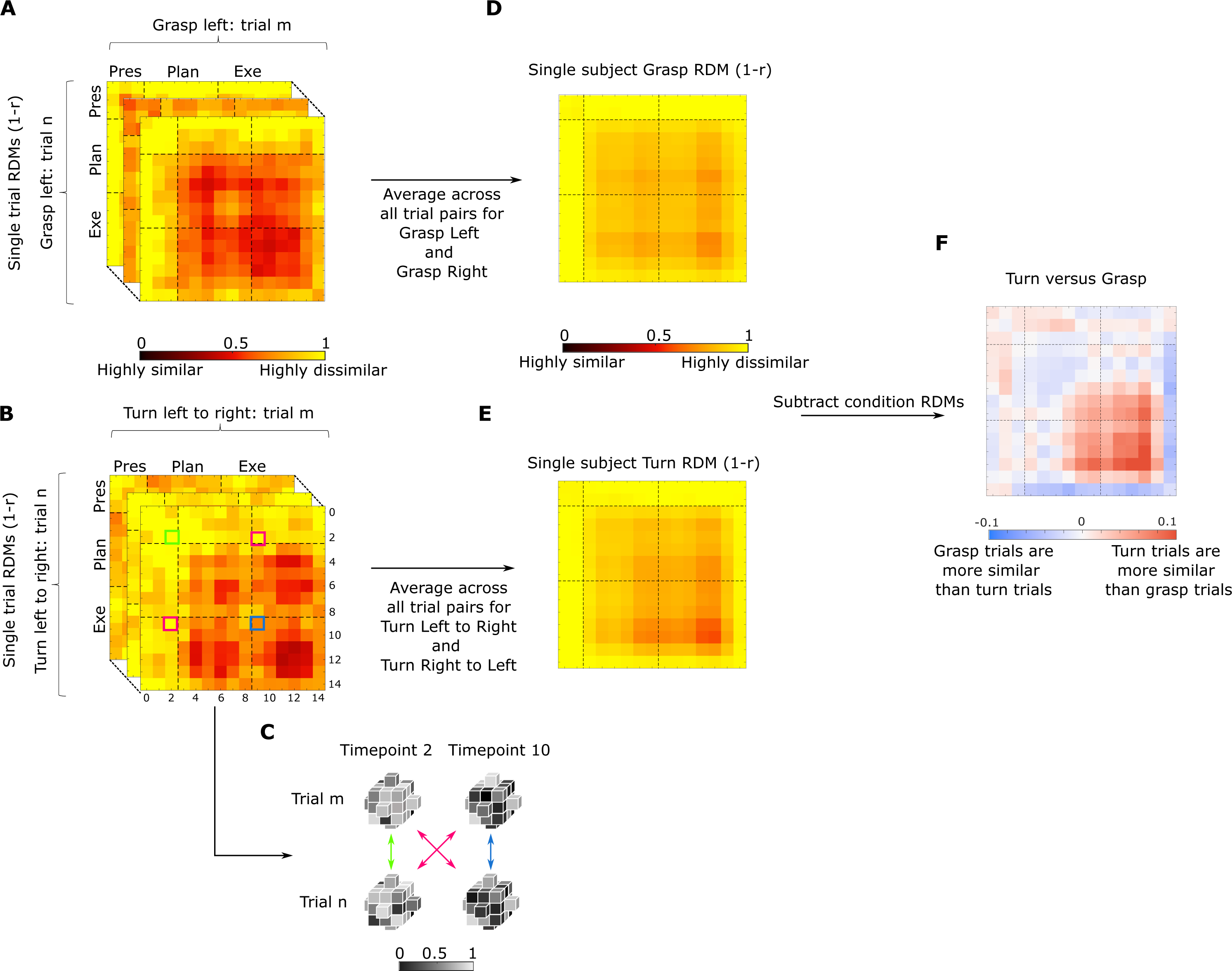
Overview of the steps to perform temporal multivoxel pattern analysis (see methods for full explanation). The figure represents the steps taken for each participant separately. Within each condition single trial representational distance matrices (RDMs) are calculated for each timepoint of each trial pairing (**3A** shows example for grasping trials and **3B** for turning trials). **3C:** Grey cubes represent voxels of the same ROI during different timepoints for two sample trials to exemplify how cells for each RDM are calculated. After calculating single trial RDMs, a condition average is calculated (middle column; **3D** and **3E** for grasping and turning respectively) which are then subtracted from each other (last column; **3F**; red and blue matrix). Not shown on picture (for results see Figure 4 and 5): the subtraction matrices are then used for bootstrapping to determine whether the group average differs significantly from zero and to investigate whether one condition is more similar than the other for each given timepoint

To further clarify calculation of the dissimilarity metric: in Figure 3B and Figure 3C, the green highlighted square indicates the dissimilarity between all the voxels (of a given ROI) at timepoint 2 of Turn Left to Right trial m with the activation of the same voxels at timepoint 2 of Turn Left to Right trial n. The magenta highlighted squares indicate the averaged dissimilarity between timepoints 2 and 10 of trial m and n. As explained before, we averaged the dissimilarity between timepoint 2 of trial m and timepoint 10 of trial n (magenta highlighted square below diagonal) with the dissimilarity between timepoint 10 of trial m and timepoint 2 of trial n (magenta highlighted square above diagonal). Finally, the blue highlighted square indicates the dissimilarity between timepoints 10 of trial m and n. In sum, values on the main diagonal show dissimilarity between within-condition trials at the same timepoint. Values that are off the main diagonal show the dissimilarity between within-condition trials at different (i.e., earlier or later) timepoints.

stRDMs were calculated for the four conditions separately, Fisher *z*-transformed and then averaged (10 % trimmed mean; Vizioli et al., 2018) within conditions (e.g., average RDM for grasp left). Finally, the averaged RDMs were then averaged across orientations (e.g., averaging of RDMs between left and right start orientations to produce average Grasp and Turn RDMs; Figure 3D and 3E respectively). This was done as we were primarily interested in investigating differences in neural representations between singular (grasping) and sequential actions (grasping then turning) and we did not expect that these differences would depend upon the start orientation. As such, this approach resulted in two RDMs for each participant.

In line Vizioli et al. (2018), we performed statistical analysis on the Fisher *z*-transformed data; however, we used the non-transformed data for visualization purposes to render the values visually more interpretable. To test for statistically significant differences between the grasp and turn RDMs, we subtracted for each participant the turn RDM from the grasp RDM resulting in a subtraction matrix (Turn versus Grasp: Figure 3F) and investigated where the subtraction differed significantly from zero. As such, a given value in the subtraction matrix that is positive indicates that grasping trials are more dissimilar than turning trials at that given timepoint. Conversely, a value that is negative indicates that turning trials are more dissimilar than grasping trials at that given timepoint. Note that we decided to statistically test whether each cell differed from zero instead of the increasingly larger sliding window analysis used in Ramon et al. (2015) and Vizioli et al. (2018). Our rationale was that the earlier studies relied on a visual task (face recognition) whereas our study relied on a motor task. Arguably, during motor preparation/execution, neural representations might evolve differently than purely visual responses. As such, we argued that statistically testing each cell separately might reveal more information on the temporal evolution of motor execution as, for instance, transitions between activation patterns between planning and execution might be brief or abrupt. Note that because we had averaged values across the main diagonal when calculating the dissimilarity matrices (Figure 3 and 5), these matrices were symmetrical reflections across the diagonal. However, we performed statistical analyses only on the values on and above the main diagonal of the Turn versus Grasp subtraction matrices (Figure 3F).

**Figure 4.**
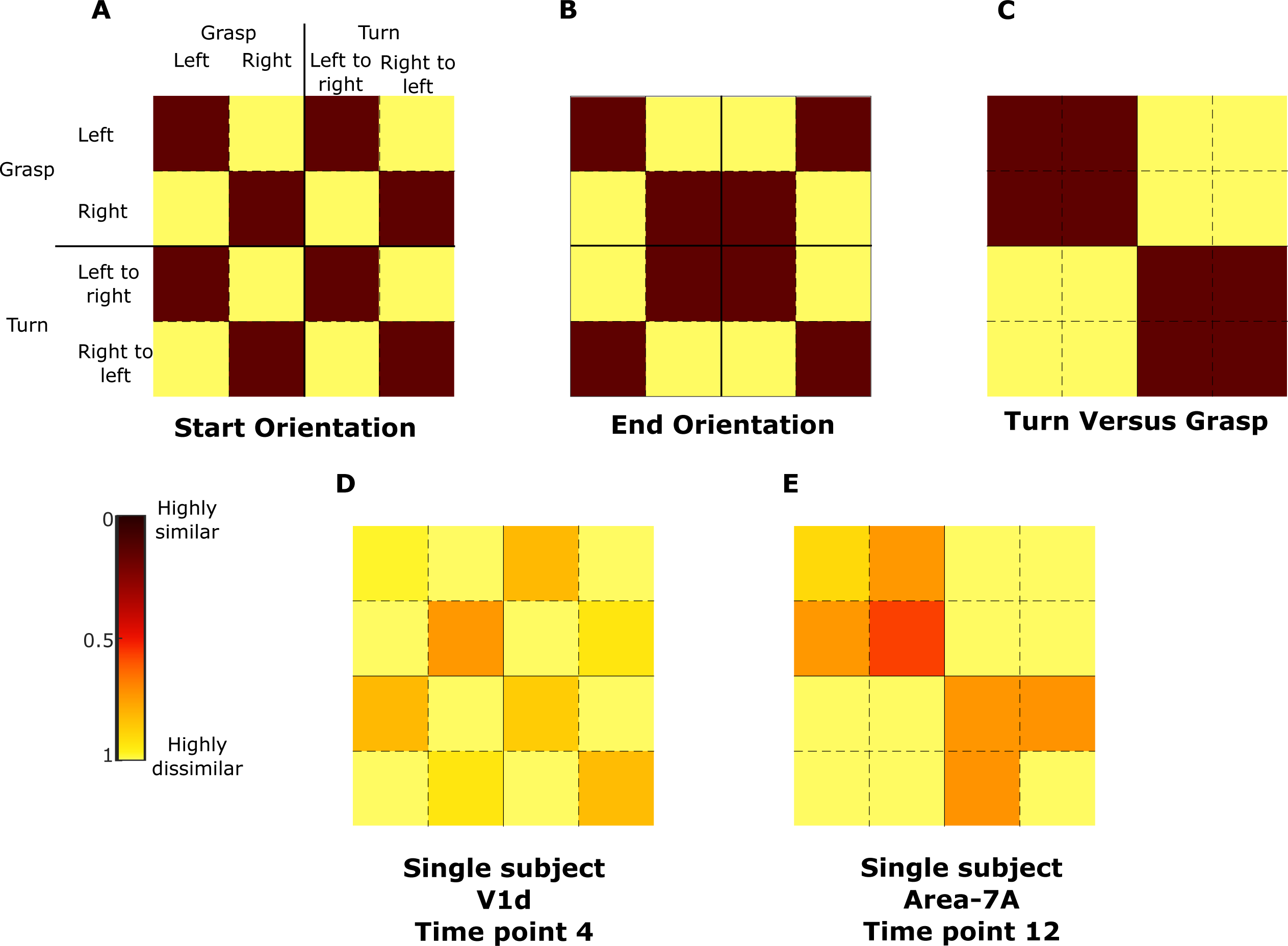
Models used for representational similarity analysis (RSA). The first row shows the models for start orientation (**4A**), end orientation (**4B**) and turn versus grasp (**4C**). The second row shows example data for one single timepoint for two regions of interest (**4D**: V1d); **4E**: Area-7A) of one participant.

**Figure 5.**
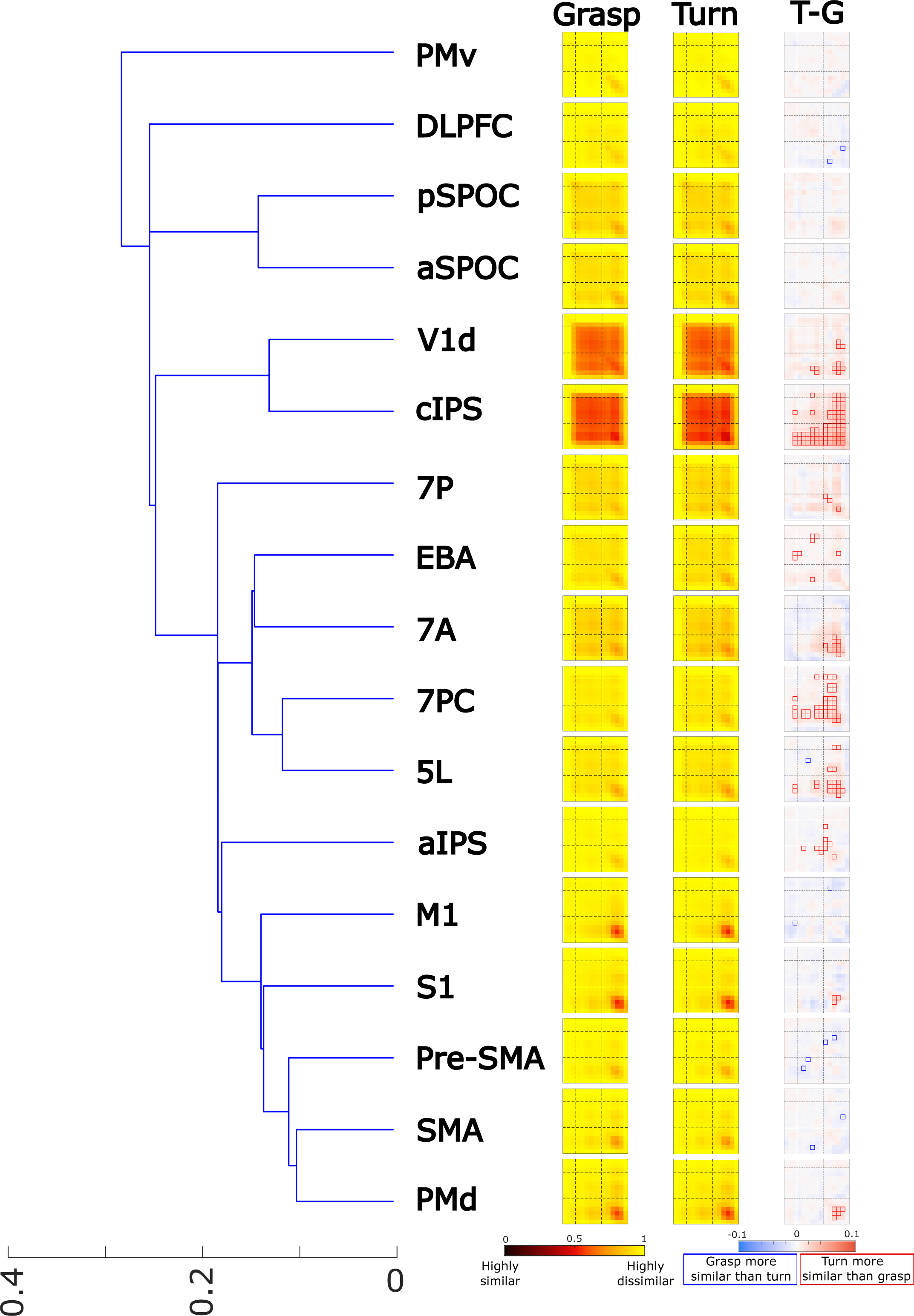
**Left:** Hierarchical clustering of all the ROIs based on the averaged representational dissimilarity matrices (RDMs) of the **Grasp** and **Turn** condition. We refer the reader to the legend of Figure 3 for the full explanation of the heatmaps. Note that **T-G** represents ‘Turn minus Grasp’ and represents the final tMVPA result showing statistically significant similarities (small red boxes = Turn > Grasp; small blue boxes = Grasp > Turn). The outline of the significance box indicates significance level. When the outline is dashed, the square was significant only when Bonferroni corrected for within-ROI comparisons. When the outline is solid, the square was significant when Bonferroni corrected for all comparisons across all ROIs. Note that we only tested whether boxes were significantly different from zero (and not from each other).

To test for statistical significance, we performed (1 - alpha) bootstrap confidence-interval analysis (critical alpha = 0.05) by sampling participants with replacement 500 times. As described above, we investigated whether tMVPA differed significantly from zero in either direction. Importantly, we excluded the first two timepoints (timepoints 0 and 1) and the last one (timepoint 14) from the statistical analyses (resulting in 12 timepoints included in the analysis: timepoints 2 to 13) because no differences were expected before the BOLD response to emerge at the start, because later timepoints reflect the post-stimulus undershoot phase of the BOLD response, and to reduce the number of comparisons requiring Bonferroni correction for multiple comparisons. See Figure 1 for the trial timing and Figure 3 for the timepoints in the tMVPA analysis).

As for the univariate analysis, we accounted for multiple comparisons using Bonferroni correction in two manners. Our first, standard, correction did not take multiple comparisons between regions in account but only the number of comparisons within an ROI. This resulted in correcting for 78 tests: the binomial coefficient of 12 timepoints (which excludes the diagonal) plus the 12 included timepoints on the diagonal. The second, strict, correction, included multiple comparisons between all regions at all time points and resulted in 936 tests. Accordingly, the critical alpha was Bonferroni corrected for the number of tests in the bootstrap confidence-interval analysis. Due to the impracticability of reporting our tMVPA data in text or table format, we provide these results only in figure format (see data availability statement). In Figure 6, cells that differ significantly from zero have a dashed outline or solid outline for the standard and strict correction, respectively.

**Figure 6.**
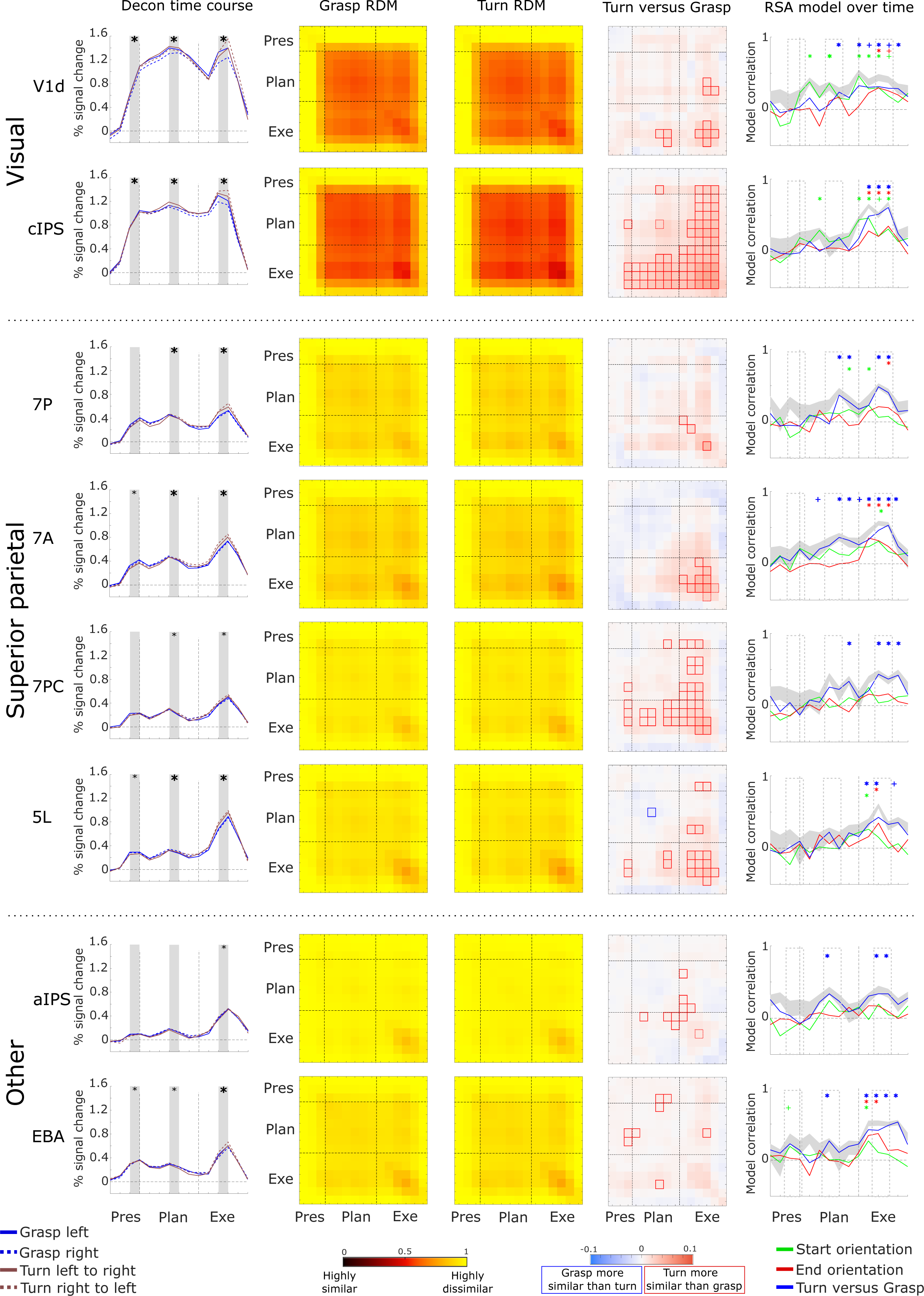
Selection of ROIs that showed prominent differences for the tMVPA analysis. Some data are reproduced from subsets of Figures 2 (Column 1) and 5 (Columns 2-4, at higher resolution) to facilitate inspection and comparisons across analysis methods for key regions. **Column 1** shows the group average deconvolution time courses for the four conditions. The X-axis represents time (in 2 s TRs) and the Y-axis represents % signal change in the BOLD signal. Vertical dashed lines represent onset of the planning phase (TR 3) (after the initial presentation phase) and the execution phase (TR 9) following the planning phase. **Columns 2 and 3** show the group average condition representational dissimilarity matrices for the grasp (column 2) and turn condition (column 3). Each cell/square of the matrix represents the dissimilarity between trials of the same condition between two timepoints. Dashed horizontal and vertical lines represent the same timepoints as those in Column 1, i.e., the first dashed horizontal/vertical line represents the start of the planning phase (TR 3) and the second dashed vertical/horizontal line represents the start of the execution phase (TR 9). **Column 4**. tMVPA results. Dashed lines have the same representation as explained for columns 1-3. Red squares indicate dissimilarity metrics between timepoints where turning trials are significantly more similar (less dissimilar) than grasping trials. blue squares represent the opposite (grasping more similar than turning). **Column 5.** Representational similarity analysis using three models, as explained in the color legend, for each timepoint of the experimental trials. Solid lines represent the correlation between the data and the model over time. The shaded grey area represents the noise ceiling with the lower and upper edges representing the lower and upper bounds. Dashed vertical lines have the same representation as explained for the previous columns. Colored “plus signs” indicate time points with statistically significant correlations for the respective model when FDR corrected for all within-ROI comparisons against zero. Colored asterisks indicate the same when FDR corrected for all comparisons (across all ROIs) against zero.

#### 2.10.3 Hierarchical clustering

The tMVPA matrices (stRDMs) for Grasp and Turn revealed that different ROIs showed different temporal profiles. For example, some regions showed similar activation patterns throughout the entire trial; whereas, other regions showed similar patterns only during motor execution (See Figure 5). To make it easier to group these different ROI timing patterns for display and discussion, we used hierarchical clustering. To do so, we averaged within-participants the grasp and turn RDMs to generate one RDM per participant. Next, we calculated for each participant the Spearman correlations between all ROIs using the averaged RDMs. The ROI correlation matrix for each participant was then transformed into a dissimilarity measure (1-*r*). Finally, hierarchical clustering was then performed using the ROI dissimilarity matrix of all participants.

#### 2.10.4 Representational similarity analysis

We used RSA to examine the degree to which the pattern of activation across voxels within each ROI at each time point represented the start orientation, the end (goal) orientation, and the task, as illustrated in Figure 4A-C. For RSA, we relied on the methods described in Kriegeskorte (2008). By examining the degree to which different conditions evoke a similar pattern of brain activation within a ROI, the nature of neural coding (or representational geometry; Kriegeskorte & Kievit, 2013) can be assessed.

For each voxel of each ROI of each participant, we performed a deconvolution GLM. This was done for each condition and each run (but not trial as for tMVPA) separately, providing the average development of brain activation over time for each condition in each voxel in each ROI of each run of each participant. Then for each voxel separately, activations of the four conditions were normalized by subtracting the grand mean (i.e., average across conditions) from the value of each condition in that voxel (Haxby et al., 2001). Note that this was done for each run separately. These steps resulted in the normalized brain activation over time averaged across trials of the same conditions within one run. This was done for each condition for each voxel of each ROI for each run of each participant.

Subsequently, we computed representational dissimilarity matrices (RDMs) within each ROI. Pearson correlations were calculated at each timepoint between all voxels of a given ROI, i.e., “within timepoint correlations”, and then transformed into dissimilarity metric (*1-r*). This was done within-condition (e.g., grasp left activations at timepoint 1 correlated with itself at timepoint 1) and between conditions (e.g., grasp left activations at timepoint 1 correlated with turn left to right at timepoint 1). In this manner, RDMs can quantify the dissimilarity in activation patterns between conditions, i.e. “how well is activity at any timepoint in a given set of voxels in a given condition correlated with the activity at the same timepoint of the same voxels during a different condition.”. Please note that we did not perform autocorrelations (e.g., correlate grasp left activations during run 1 with itself thus leading to correlation values of 1). Instead to test within-subject reliability of the RDM calculations, we used cross-validation by splitting data into all potential combinations of runs yielding two sets of four runs each (two examples: split 1 = correlate all even runs with the uneven runs; split 2: correlate the first four runs with the last four runs). Split data provides data-based estimates of dissimilarity even for the same condition as no autocorrelations are performed whereas, in unsplit data, the dissimilarity is necessarily zero. Data for all cells in the RDM is necessary for RSA on factorial designs to ensure that contrasts are balanced across orthogonal factors. Finally, the RDMs were Fisher transformed to have a similarity metric with a Gaussian distribution.

To test whether each region contained information about condition differences, we measured correlations between the RDM in each region (Figure 4D-E) and three separate models that capture orthogonal components of the experimental task (Figure 4A-C). This was done for each timepoint separately (15 timepoints included in the GLM being timepoint 0-14; time-resolved decoding). Note that in contrast to the tMVPA we included all timepoints due to our selection of models which included “two visual models” which may decode already during the earliest timepoints. In total, we used three models that capture specific task attributes: (A) Start orientation, irrespective of task (i.e., grasp or turning) — a timepoint with an RDM that correlates with this model would indicate an encoding of the initial orientation of the handle (i.e., left, or right), (B) end orientation, irrespective of the performed action (i.e., grasp or turning) — a timepoint with an RDM that correlates with this model would indicate an encoding of the final orientation of the handle (i.e., left, or right) after task execution, (C) motor task, irrespective of the initial/final handle’s orientation (i.e., left, or right) — a timepoint with an RDM that correlates with this model would indicate an encoding of the task goal (i.e., grasping or turning). The metric was calculated by computing the Spearman correlation (ρ) between the Fisher-transformed split RDMs and each model for each ROI for each participant. For each ROI we also calculated the upper and lower bound of the noise ceiling. Briefly, the noise ceiling is the expected RDM correlation achieved by the (unknown) true model, given the noise of the data and provides an estimate of the maximum correlations that could be expected for a given model in a given ROI (Kriegeskorte, 2008). RDMs were first rank transformed, following the original *z*-transformation. The upper bound of the noise ceiling, considered an overestimate of the maximum correlation, was calculated as iteratively correlating one participant’s RDM with the average RDM of all participants (thus including the given participant) and then averaging across all participants. The lower bound of the noise ceiling, considered an underestimate of the maximum correlation, was calculated as iteratively correlating between one participant’s RDM with the average RDM of all other participants (thus excluding the given participant), and then averaging across participants.

We used one-way Student’s *t* tests (alpha < 0.05) to assess whether model correlations were significantly greater than zero for each timepoint in each ROI. Importantly, as for the tMVPA we decided to exclude the first two timepoints (timepoints 0 and 1) and the last one (timepoint 15). This resulted in 13 timepoints included in the analysis. Note that we did not compare whether models outperformed each other as we were primarily interested in whether which ROI embodied what type of information. To be in line with previous studies combining motor tasks and RSA (e.g., Di Bono et al., 2015; Gallivan et al., 2013; Monaco et al., 2020), we performed corrections for multiple comparisons using the false discovery rate. As for the univariate analysis, we performed multiple comparisons for each ROI separately (i.e., 12 timepoints x 3 models) and in a stricter manner across all regions (strict comparison: 12 timepoints x 3 models x 17 ROIs). The results can be found in Figure 6: models at any given timepoint that are significantly larger than zero for only the within-ROI corrections are indicated with a “plus sign”. When the model at the timepoint also reaches significance for the strict threshold, this is indicated by an asterisk instead. Due to the amount of data (i.e., 12 timepoints x 3 models x 17 ROIs), we provide a specific selection of values in the results section. Akin to the univariate analysis, we provide data for timepoints 2 and 3 for the presentation phase, 6 and 7 for the planning phase and 11 and 12 for the execution phase in Table 2 for the subset of ROIs in Figure 6. Again, note that we performed statistical tests and performed multiple comparisons for 12 timepoints and not just the six timepoints mentioned here to keep the statistical analysis in line with our tMVPA approach. In addition, note that here we do not average timepoints for each phase (e.g., no averaging of timepoint 2 and 3 into one “presentation phase”) for the same reasons. The values in Table 2 reflect the *q* values resulting from the strict false discovery rate correction as well as the confidence intervals, given as [lower threshold mean upper threshold], which are Bonferroni corrected for multiple comparisons in the strict manner (12 timepoints x 3 models x 17 ROIs). Note that we had to correct confidence intervals using Bonferroni given the step-by-step approach FDR, which is not common for confidence intervals. As such, it is possible that our actual interpretation of the statistical analysis using FDR (shown in Figure 6) reveals significant effects despite the confidence intervals using Bonferroni (shown in Table 2) containing zero.

**Table 1.** The average number of functional (1-mm iso-)voxels per Region of Interest. Values shown are mean ± SEM.

| Region of Interest | Number of Functional Voxels |
| --- | --- |
| M1 | 98 ± 7 |
| S1 | 123 ± 11 |
| PMv | 107 ± 8 |
| PMd | 217 ± 9 |
| aSPOC | 178 ± 14 |
| pSPOC | 138 ± 7 |
| aIPS | 268 ± 17 |
| cIPS | 172 ± 10 |
| Pre-SMA | 108 ± 7 |
| SMA | 132 ± 10 |
| DLPFC | 129 ± 9 |
| V1d | 152 ± 13 |
| Area-5L | 167 ± 10 |
| Area-7A | 194 ± 12 |
| Area-7P | 150 ± 9 |
| Area-7PC | 198 ± 13 |
| EBA | 1163 ± 23 |

**Table 2.**
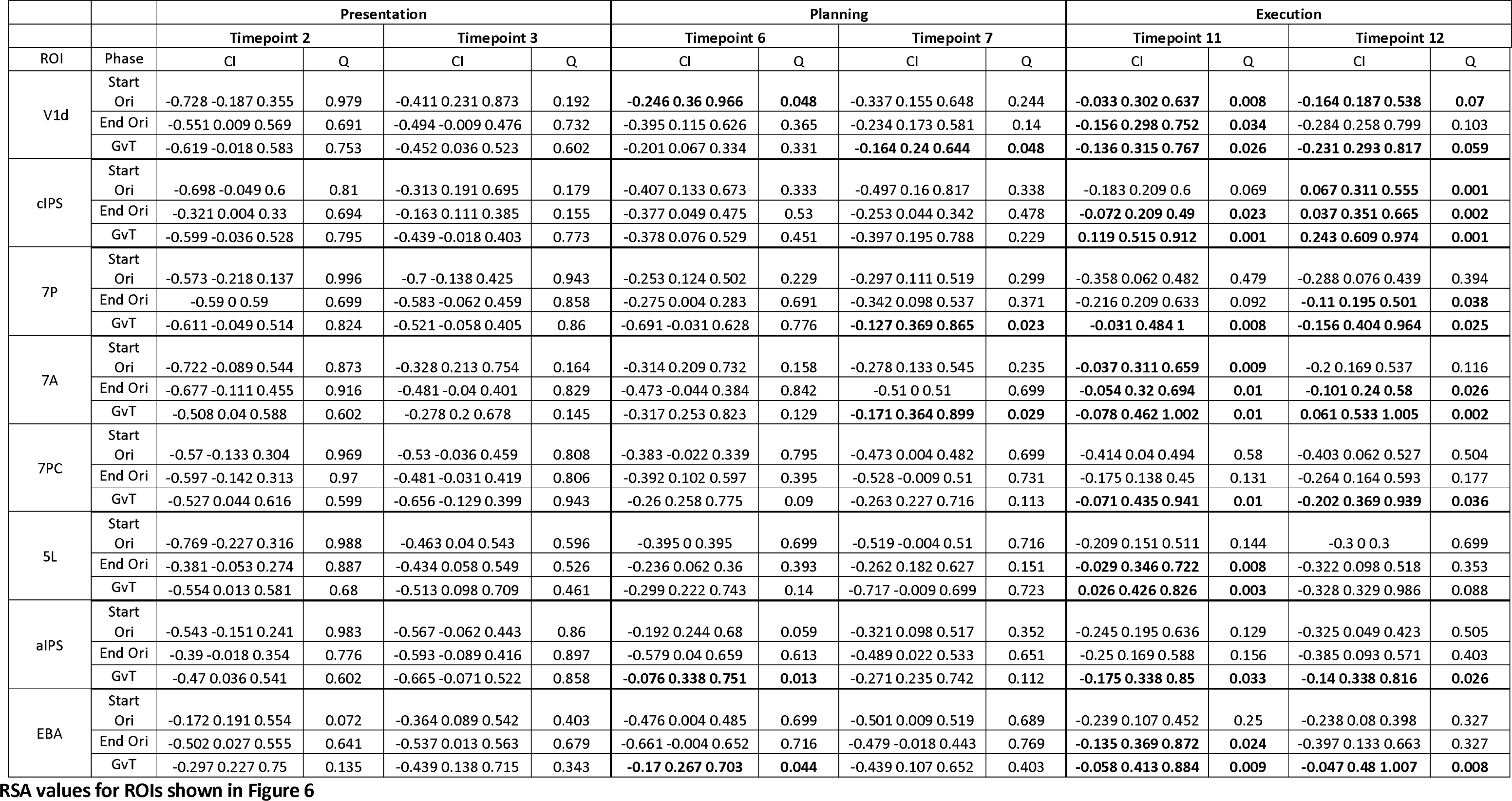
Values represent model correlation values for each ROI at specific timepoints. Values depict mean ± confidence interval and is written as [lower threshold mean upper threshold]. Note that the confidence intervals are Bonferroni corrected in the strict manner (critical alpha divided by 12 timepoints x 3 models x 17 ROIs). Q-statistic represents the p-value that is FDR corrected in the same strict manner. Significance was defined as a Q-value < 0.05 (see 2.10.4 Representational Similarity Analysis).

To end, we will also qualitatively discuss the grasp and turn RDMs (which subtraction constitutes the tMVPA subtraction matrix that is statistically tested) in order to facilitate conceptual understanding of the data. Please note that this qualitative discussion will not be incorporated in the results and discussion due to the lack of the statistical analysis (which has been done on the subtraction matrix instead).

## 3. Results

Qualitative examination of the deconvolution time courses (Figure 2) and the tMVPA results indicated that different regions had different temporal profiles of activity. As shown in Figure 5, hierarchical clustering of the RDMs averaged across Grasp and Turn facilitated conceptual grouping of the ROIs for further investigation.

Notably, ROIs differed in the time ranges over which they showed reliable activation patterns across trials of the same type. Most strikingly, V1d and cIPS showed strong temporal similarity in activity patterns throughout the Plan and Execute phases of the trial, as indicated by the large block of high similarity (dark red) beginning early in planning and continuing through late execution (Figure 5). That is, in these regions, voxel activation patterns were not only similar during the same phase (along the diagonal cells) but were also similar across timepoints throughout planning and execution (off-diagonal cells in the planning and execution phases). This indicates consistency in the neural representation throughout the trials. Similar but weaker similarity patterns were also observed in other regions, particularly aSPOC, pSPOC, 7P, EBA, 7A, 7PC, and 5L.

In contrast, other regions -- aIPS, M1, S1, pre-SMA, SMA, PMd -- showed trial-consistent activation patterns predominantly for the peak execution period (dark red blocks in the execution phase; i.e., in lower right corners of Grasp and Turn matrices). In some cases, there was also some consistency of patterns between the peak execution phase and earlier timepoints during Plan and Execute of which M1 is a clear example in Figure 5 (off-diagonal cells between planning and execution phase, as indicated by the reddish “wings” above and left of the peak similarity). PMv and DLPFC showed only weak consistency of patterns across trials, highest in the peak of execution.

Interesting patterns were also observed in the tMVPA difference in trial-by-trial consistency between turn and grasp (final column of Figure 5). Notably many of the regions showed higher consistency during Turn actions than Grasp actions.

Based on these observations from Figure 5, we focussed subsequent analysis and interpretation on a subset of ROIs that is shown in Figure 6.

### 3.1 Visual regions of V1d and cIPS

#### Univariate analysis

The first column in Figure 6 shows the deconvolution time courses for V1d and cIPS. V1d and cIPS show a steep increase in average voxel activation following object presentation (V1d = 0.31 ± 0.06, P < 0.001,P(strict) = 0.008; cIPS = 0.46 ± 0.07, P < 0.001; P(strict) < 0.006 compared to baseline. This initial increase is maintained throughout the planning (V1d = 1.31 ± 0.19, P < 0.001,P(strict) < 0.001; cIPS = 1.09 ± 0.19, P < 0.001; P(strict) < 0.001) and execution phases (V1d = 1.07 ± 0.22, P < 0.001,P(strict) = 0.013; cIPS = 1.15 ± 0.18, P < 0.001; P(strict) < 0.001).

#### Qualitative (non-statistical) assessment of the condition RDMs

In Figure 6 the second and third columns show the condition RDMs for grasp and turn, respectively. For both regions, activation pattern similarity within conditions strongly increases following object presentation. Due to the sluggish nature of the hemodynamic response and our findings for the univariate analysis, it is plausible that this early increase of similarity in the activation patterns is primarily driven by a visual response. Notably, the cIPS within-condition similarity in activation patterns increases towards the peak execution (approx. timepoint 11-13).

#### Temporal multivoxel pattern analysis

The fourth column in Figure 6 depicts the subtraction Turn minus grasp on which we performed statistical analyses as previously described. For V1d, tMVPA is unable to discriminate the grasp and turning condition for matched timepoints (along the diagonal) during motor planning. That is, subtracting the grasp dissimilarity from the turn dissimilarity (second and third column in Figure 6) does not result in a difference that differs statistically from zero. Interestingly, tMVPA is able to discriminate the activation patterns of grasping and turning during late execution (i.e., the “plus sign” of significant squares in the grasp versus turn RDM for V1d) which indicates that V1d may encode change in visual information (e.g., increased motion during) or motor-components that are non-visual. Interestingly, tMVPA was also able to discriminate activity patterns related to grasping versus turning when dissimilarity calculations were done between timepoints of the late planning phase and timepoints of the late execution phase (i.e., the three adjacent significant squares in the grasp versus turn RDM for V1d). These findings suggest that the activation patterns in V1d become tuned towards either grasping or turning during late motor planning and may not be limited to solely encoding visual information. Similar but stronger effects were found for cIPS. tMVPA was able to discriminate cIPS activation patterns during most of the execution phase as well as when dissimilarity calculations were done between timepoints of the execution phase and early and late timepoints of the planning phase (i.e., the vertical and horizontal “wings” of significant squares above and left of the execution phase). This suggests that in cIPS, representations of the task emerge early in the planning period.

#### Representational similarity analysis

For the RSA, we tested how well multivariate activation patterns fit three models for the representation of start orientation, end orientation, and task (Turn versus Grasp) over time (Figure 6; last column). The confidence intervals and *q-*values for a selection of timepoints (Figure 6; last column; timepoints within transparent rectangles with dashed grey outlines) is provided in Table 2. Notably, although all three types of information appear to be represented in both V1d and cIPS during action execution, starting orientation appeared to be represented during the early planning phase. In addition, during the late planning phase, both starting orientation and grasp versus turn also appeared to be represented in V1d (see Table 2 for values for Timepoints 6 and 7).Although the decoding of start orientation during planning and end orientation during execution may reflect the processing of simple visual orientation information (Kamitani & Tong, 2005), coding of both start and end orientation overlapped during execution (Figure 6; last column; V1d and cIPS; green and red trace), suggest a much more complex representation. Notably, V1d, represented the task during planning, before any action had been initiated, perhaps due to anticipation of the visual consequences of the upcoming action.

In sum, while our findings are consistent with a role for V1d and cIPS as predominantly visual regions, the similarity in activation patterns (tMVPA) between the planning and the execution phase, in particular for cIPS (Figure 6; fourth column) as well as V1d coding the motor task during planning (Figure 6; last column’ blue trace) suggest that these regions may encode motor components already during the planning phase.

### 3.2 Regions of the SPL: 7A, 7PC, 5L and 7P

#### Univariate analysis

Despite only 7A showing a significant increase following object presentation (area 7P = 0.13 ± 0.05, P = 0.09, P(strict) = 1.00; area 7A = 0.16 ± 0.05, P = 0.037, P(strict) = 0.63; area 7PC = 0.11 ± 0.06, P = 0.36, P(strict) = 1.00; area 5L = 0.15 ± 0.05, 0.040. P(strict) = 0.679), all regions showed a significant increase following action planning (area 7P = 0.40 ± 0.08, P < 0.001, P(strict) < 0.001; area 7A = 0.42 ± 0.08, P < 0.001, P(strict) = 0.01; area 7PC = 0.25 ± 0.07, P = 0.006, P(strict) = 0.103; area 5L = 0.28 ± 0.06, P = 0.001, P(strict) = 0.017), and execution (area 7P = 0.36 ± 0.07, P < 0.001, P(strict) = 0.009; area 7A = 0.49 ± 0.09, P < 0.001, P(strict) = 0.003; area 7PC = 0.30 ± 0.08, P = 0.006, P(strict) = 0.104; area 5L = 0.54 ± 0.09, P < 0.001, P(strict) = 0.002). As shown in the first column in Figure 6, given that the activity within this ROIs show a different trend than cIPS and V1d (i.e., primarily a significant increase in activation following action planning, not object presentation), it seems plausible that the subregions of the SPL have primarily a visuomotor involvement.

#### Qualitative (non-statistical) assessment of the condition RDMs

The condition RDMs of Figure 6 show that the ROIs within the SPL show activity patterns that are similar to each other. First, similarity between trials within each condition increases following object presentation, in particular, during early planning. Second, similarity in activation patterns is then maintained until a second increase during the execution phase. Notably, this pattern (in the condition RDMs of the SPL regions) is similar to that of V1d and cIPS, although it seems weaker (i.e., less red). Most importantly, the increase in within-condition trial similarity following object presentation seems weaker for the SPL regions than for V1d and cIPS suggesting that these regions are likely less modulated by this type of motor task.

#### Temporal multivoxel pattern analysis

In line with the assessments of the univariate analysis and the condition RDMs, tMVPA indicate that the regions of the SPL may be involved in the motor task, in particularly during motor execution (Figure 6; column 4). For area 7A, tMVPA was able to discriminate activation patterns for grasping and turning during the later execution phase. Interestingly, we found similar results for area 7PC as for cIPS (as explained in “Visual regions of V1d and cIPS”). tMVPA was able to discriminate grasping and turning throughout both earlier and later stages of the execution phase as well as when dissimilarity calculations were done between timepoints of the planning phase and timepoints of the execution phase. These findings indicate that area 7PC might anticipate turning actions already during the planning phase. The results for area 5L are somewhat similar to the findings for areas 7A and 7PC. That is, tMVPA is primarily able to differentiate between grasping and turning during the later execution phase but also, in a more limited manner than area 7PC, when dissimilarity was calculated on timepoints of the early/late planning phase and timepoints of the execution phase. For area 7P, tMVPA could significantly discriminate activation patterns associated with grasping and turning only during the execution phase. Finally, it can be seen that tMVPA can discriminate between grasping and turning by relying on the activation patterns of each of the four regions. However, this ability to discriminate the task is the weakest in 7P, stronger in 7A and 5L and the strongest in 7PC.

#### Representational similarity analysis

The RSA results (Figure 6 column 5) indicate that the regions of the SPL could rather have a visuomotor involvement than a purely visual one: starting orientation cannot be decoded from the SPL (Figure 6; last column; 7P, 7A, 7PC and 5L; green trace) during the planning phase. In support of our tMVPA results, grasping versus turning can be decoded from all four regions in the SPL (Figure 6; last column; 7P, 7A, 7PC and 5L; blue trace). RSA is primarily able to do this when relying on the execution phase. Significant decoding is also observed during the late planning phase for the regions 7A and 7P (see Table 2 for values for timepoints 6 and 7). Interestingly, in area 7PC only turning versus grasping can be decoded, suggesting that this region might be mainly involved in rotating the wrist, irrespective of the direction, and may not integrate visual information. Indeed, from the other regions (areas 7A, 5L and 7P) both start and end orientation can be decoded during the action execution phase suggesting that these regions may encode specific wrist orientations, or may still be involved in visuomotor integration.

In sum, both tMVPA and RSA seem to provide evidence for the involvement of the SPL in decoding sequential actions. That is, sequential (grasping then turning) actions can be differentiated from singular grasping actions based on the activation patterns from the regions 7A, 7PC, 5L and 7P. In particular, for areas 7A and 7P our findings highlight that RSA is already able to do so during motor planning.

### 3.3 EBA and aIPS

#### Univariate analysis

As shown in Figure 6, EBA shows an initial response to object presentation (EBA = 0.19 ± 0.05, P = 0.011, P(strict) = 0.184) which is maintained throughout action planning (EBA = 0.27 ± 0.07, P = 0.008, P(strict) = 0.140) and execution (EBA = 0.29 ± 0.06, P = 0.002, P(strict) = 0.037). In contrast, for aIPS the univariate analysis reveal only a significant response to action execution (aIPS = 0.24 ± 0.07, P = 0.016, P(strict) = 0.273) that is absent for the presentation (aIPS = 0.03 ± 0.06, P = 1.00, P(strict) = 1.00) and planning phase (aIPS = 0.25 ± 0.05, P = 0.07, P(strict) = 1.00).

#### Qualitative (non-statistical) assessment of the condition RDMs

In line with the univariate analysis, the condition RDMs for aIPS show that activation patterns remain very dissimilar and then show an increase following action execution. As such, the condition RDMs indicate that aIPS is primarily involved in action execution. In the condition RDMs for EBA, a clear visual response can be seen as within-condition similarity increases early during trials. Interestingly, in line with the univariate analysis, within-condition similarity increases again following task execution supporting the notion of EBA’s involvement in motor execution and/or visual feedback of the hand, which in the case of the condition RDMs would relate to the difference in perceived motion.

#### Temporal multivoxel pattern analysis

tMVPA can discriminate activation patterns in aIPS related to grasping and turning during the early execution phase, again supporting the notion that aIPS is involved in the execution of hand-object interactions. Interestingly, tMVPA was able to discriminate activation patterns related to grasping and turning during the planning phase as well as when dissimilarity was calculated between timepoints of the planning phase and timepoints of the execution phase. EBA showed sporadically significant differences between the two tasks during the early visual response and planning.

#### Representational similarity analysis

From aIPS, RSA could discriminate the activation patterns associated with grasping and turning during the planning phase as well as during the execution phase (Figure 6; last column; aIPS; blue trace; Table 2 for data of timepoints 6 and 7). For EBA, the RSA revealed an early visual response as it was able to decode starting orientation from the activation patterns following object presentation (Figure 6; last column; EBA; green trace). Interestingly, in with the previous analyses, RSA was able to decode grasp versus turn in both the planning phase as well as the execution phase (Figure 6; last column; EBA; blue trace).

In sum, our findings for aIPS are in line with previous studies that showed its involvement in motor planning and execution (e.g., Singhal et al., 2013). Interestingly, our multivariate analyses support the suggestion that EBA, perhaps along with hand-selective divisions of the lateral occipitotemporal cortex, is not a purely visual region but is also involved in analyzing visual feedback of the body and hand during motor execution (Astafiev et al., 2004; van den Heiligenberg et al., 2018; Zimmermann et al., 2016)

### 3.4 Other regions of interest

We focussed the presentation of the results above in the most interesting findings about how hand-object interactions unfold over time. Other regions showed effects that were absent, weaker or less surprising. The condition RDMs and tMVPA results for the previously discussed ROIs and other ROIs can be found in Figure 5. The RSA findings of all regions can be found on GitHub (https://github.com/GuyRens/OvenDials).

Briefly, we found weak effects (no significant tMVPA) for the parietal regions of pSPOC and aSPOC (tMVPA: no significant squares; RSA: no significant models) which have been considered to be typically involved in hand-object interactions. For DLPFC the tMVPA revealed only one significant difference for grasp vs turn during late execution. RSA revealed that both the end orientation and motor task model were significant for at least one timepoint (when applying strict FDR, i.e., taking all ROIs in account when correcting for multiple corrections) indicating the involvement of DLPFC in motor execution. For SMA and pre-SMA, the RSA revealed that the start orientation model was significant for SMA at timepoint 4 and the end orientation model at timepoint 10 for pre-SMA. Both regions showed significant negative values (grasp > turn) for limited cells when correlations were computed on the planning and execution phase. For S1, RSA showed significance for the start orientation and motor task during late execution. This is supported by tMVPA results which showed higher similarity for turn than for grasp during this phase. Finally, both PMv and M1 show no response in the tMVPA results. It should be noted in Figure 5 that the subtraction matrix shows a significant cell for M1 being a correlation between the presentation phase and the execution phase. Similarly, for the RSA results the end orientation model is significant PMv timepoint 3. Both these results are likely to be analytical anomalies as participants had no information about the upcoming motor task at these stages of the experimental trials. To end, both PMv and M1 reached significance for the grasp versus turn model for one timepoint during motor execution further supporting the well-accepted of these regions involvement in motor control.

## 4. Discussion

Our results (summarized by the outlined circles in Figure 2) provide new insights into how neural representations unfold over time for a simple prehension task vs. a more complex manipulation task. While prior work has shown that different types of grasp sequences can be decoded (Gallivan et al., 2016), the current study examines the temporal coding of this information across planning and execution, as well as the nature of the information being represented in individual brain regions over time. In addition, our study examines the activity of regions within the SPL, not previously explored in the prior work. Our additions corroborate earlier findings (Gallivan et al., 2016, 2019) that action sequences are represented in visual areas (such as V1 and EBA), but also implicate functional subdivisions of the SPL (areas 7A, 7PC, 7P and 5L).

Using RSA, we found that while some regions, particularly V1, coded start orientation during planning, multiple regions (V1, cIPS, SPL and EBA) coded a combination of start orientation, end orientation and task during execution. This finding suggests that the activation patterns in these areas reflect more than simple visual-perceptual information such, as the visual orientation of the dial (Kamitani & Tong, 2005); instead, these patterns reflect the combination of the initial goal, the motor specific act, and the final outcome. Moreover, the similarity of tMPVA activation patterns across planning and execution, particularly in cIPS and area 7PC, suggests that an action and its outcome are anticipated well before the movement begins.

### Temporal Unfolding of Complex vs. Simple Actions

We investigated the temporal unfolding of complex vs. simple actions using multiple measures of neural responses: univariate time courses (Figure 1), RSA for sequential volumes (Figures 4 and 6) and, tMVPA (Figures 3, 5 and 6). These different measures provide a fuller picture of how actions unfold over time than considering each approach in isolation. For example, univariate signals reveal that both V1 and cIPS show robust visual signals in all three phases of the trial (presentation, plan and execute), as do SPL regions and EBA to a lesser degree. tMVPA shows consistency in the patterns of activation in these areas throughout the trial. However, RSA shows that different areas are representing different kinds of information at different points in the trial. For example, V1 represents start orientation early in the trial and task later, while 7PC represents task primarily starting from task onset. Moreover, tMVPA shows that trial-by-trial similarity during execution and between execution and planning is higher for Turn than Grasp trials.

These results build upon earlier work examining univariate and multivariate time courses during motor tasks (Ariani et al., 2018b; Gallivan et al., 2016c). They also provide a valuable complement for EEG studies that reveal temporal coding with even finer temporal resolution but poorer spatial resolution. An EEG study by (Guo et al., 2019) showed that grasp orientation (defined by instruction rather than object attributes) can be classified during both a visual preview and action execution, with similar representations between the two phases; moreover, their most informative electrodes were over left caudal parietal cortex, though source localization indicated diverse potential sources (that may include cIPS, SPL, and EBA).

One of the more surprising results of our TMVPA and RSA findings was the involvement of SPL in showing differences between Turn and Grasp actions, in particular areas 7A and 7PC. The role of the SPL in hand actions has been relatively less studied than areas in the IPS (especially cIPS and aIPS) and SPOC (V6/V6A), for which clear homologies between humans and other primates have been proposed (Culham & Kanwisher, 2001; Grefkes & Fink, 2005). One partial exception is area 7PC, an SPL region dorsal to aIPS, that has been proposed as the human homologue of the macaque medial intraparietal sulcus (mIPS) and/or parietal reach region (PRR), albeit with limited consensus about locus (Gallivan & Culham, 2015) and putative homologies (Culham et al., 2006). These SPL areas likely form part of the dorsomedial (or dorsal-dorsal) visual stream (Rizzolatti & Matelli, 2003) and have been postulated as an integration zone for visual, motor, and somatosensory information for goal-directed movements (Passarelli et al., 2021). Human and macaque SPL have similar structural organization with the anterior sector -- 7PC (macaque homologue: VIP; Caminiti et al., 2015) and 5L (macaque homologue: PE; Gamberini et al., 2020) -- being predominantly somatosensory and the more caudal parts -- 7A (macaque homologue: PEc; Gamberini et al., 2020) and 7P (macaque homologue: V6A; Gamberini et al., 2020) -- being both somatosensory and visual (Gamberini et al., 2020). The anterior regions of the SPL (5L and 7PC) fall on the more somatosensory side of the visual-to-somatosensory gradient from the posterior to anterior SPL. As such, they may be involved in anticipating the sensory (perhaps largely somatosensory) and motor requirements and consequences of the action.

### Orientation Processing

Our experiment explicitly involved tasks that required processing of object orientation for grasping and turning. While processing of object location for reaching and object shape and size for grasping have been well characterized behaviorally and neurally (Cavina-Pratesi et al., 2010), the influence of object orientation on reach-to-grasp actions has only recently been investigated in the monkey and human. These studies have suggested that object orientation and/or wrist/grip orientation rely on V6A/aSPOC (Battaglini et al., 2002; Monaco et al., 2011) and the adjacent cIPS (Rice et al., 2007; Shikata et al., 2001; Valyear et al., 2006) perhaps along with aIPS (Breveglieri et al., 2022). One fMRI study that had participants grasp a dial much like ours found that “bistable” object orientations (which equally afforded two possible grip postures) evoked greater activation in cIPS than stable orientations (which afforded only one grip posture) (Wood et al., 2017). Moreover, this study found that two neuropsychological patients with unilateral damage to cIPS showed deficits in grasping bistable orientations with the contralateral hand. We find that cIPS codes a combination of start orientation, end orientation, and task during action execution. Moreover, cIPS activation is more consistent throughout a trial in a complex turning task, where object orientation must be changed through a grip rotation, than a simpler grasp-only task, corroborating the proposed role of cIPS in processing orientation for action. To our surprise, we did not observe any orientation coding in aSPOC or pSPOC.

Akin to cIPS, early visual cortex has been increasingly implicated in motor planning and execution (Gallivan et al., 2014, 2019; Gutteling et al., 2015; Monaco et al., 2020). Our findings suggest that the visual cortex (V1) codes the start orientation of the object throughout the trial with increases in the task representation toward the end of planning and during execution. This suggests that V1 activation patterns are not solely driven in a bottom-up fashion by current visual cues (Kamitani & Tong, 2005) but come to incorporate the complexity of the task and visual information as the action unfolds.

Although studies of EBA have focused on its role in body perception (Kontaris et al., 2009), EBA has also been implicated in processing body posture during planning and execution of goal-directed actions (Gallivan et al., 2016; Zimmermann et al., 2018). We found some evidence for this notion: EBA has strong representations of task and orientations during execution but only limited ones during planning. As such, EBA may be mainly driven by the perceptual consequences of the action.

### Methodological considerations

Our “grasping” actions were atypical in that they required gripping the dial between the knuckles of the index and middle fingers, in a way that was similar between grasping, turning to the left, and turning to the right. This choice of posture was deliberate, to ensure that the representations of the upcoming task were not confounded with differences in initial grip posture. Nevertheless, this strategy differs considerably from the approach that participants would likely have undertaken without such instruction (Rosenbaum et al., 1990). Nevertheless, we found multivariate differences even in the case when the low-level grasp posture remained constant (Ariani et al., 2015; Ramon et al., 2015). Finally, we focused on the dorsal pathway given the low motor complexity (Goodale & Milner, 1992). However, using more complex objects could be used to explore ventral stream involvement: Dijkerman et al, (2009) highlighted its involvement in high-level grip selection. In addition, it has been argued that the ventral stream, in particular EBA (Zimmermann et al., 2016) provides an initial structure to the motor plans for grasping whereas regions of the SPL might be more feedback-driven (Scheperjans, Hermann, et al., 2008). Techniques with high temporal resolution such as TMS and EEG could explore the relationship between these regions that have similar activation patterns in our study (Figure 6) but that have been considered to be functionally different.

## Conclusion

We investigated the neural underpinnings of how hand-object interactions unfold on a moment-to-moment basis. Importantly, we found that the neural underpinnings of grasping and turning actions could best be differentiated in occipital cortex (particularly cIPS and V1) and SPL. These results corroborate the importance of cIPS in processing object and grip orientation and suggest that SPL may play a larger role in action sequences than previously realized. Our results also show how tMVPA can be used to understand how motor actions are represented across different phases of the trial.

## Acknowledgements

This work was supported by grants from the Canadian Institutes of Health Research Operating Grant (MOP84293 to J.C.C.), Western University Strategic Success Program (J. C. C.), the Natural Sciences and Engineering Research Council of Canada (to J.C.C.) and a CIHR-Doctoral Research Award 215131 (T.D.F).

## Conflict of interest

The authors declare no conflicts of interest.

## Notes

https://github.com/GuyRens/OvenDials

### Competing Interest Statement

The authors have declared no competing interest.

### Summary of Updates

Updates to the manuscript after feedback from reviewers of the Journal of Neuroscience

## Bibliography

Ariani, G., Oosterhof, N. N., & Lingnau, A. (2018). Time-resolved decoding of planned delayed and immediate prehension movements. Cortex, 99, 330–345. https://doi.org/10.1016/j.cortex.2017.12.007

Ariani, G., Wurm, M. F., & Lingnau, A. (2015). Decoding internally and externally driven movement plans. Journal of Neuroscience, 35(42), 14160–14171. https://doi.org/10.1523/JNEUROSCI.0596-15.2015

Astafiev, S. v., Stanley, C. M., Shulman, G. L., & Corbetta, M. (2004). Extrastriate body area in human occipital cortex responds to the performance of motor actions. Nature Neuroscience, 7(5), 542–548. https://doi.org/10.1038/nn1241

Badre, D., & Nee, D. E. (2018). Frontal Cortex and the Hierarchical Control of Behavior. Trends in Cognitive Sciences, 22(2), 170–188. https://doi.org/10.1016/j.tics.2017.11.005

Battaglini, P., Muzur, A., Galletti, C., Skrap, M., Brovelli, A., & Fattori, P. (2002). Effects of lesions to area V6A in monkeys. Experimental Brain Research, 144(3), 419–422. https://doi.org/10.1007/s00221-002-1099-4

Beurze, S. M., De Lange, F. P., Toni, I., & Medendorp, W. P. (2007). Integration of target and effector information in the human brain during reach planning. Journal of Neurophysiology, 97(1), 188–199. https://doi.org/10.1152/jn.00456.2006

Beurze, S. M., De Lange, F. P., Toni, I., & Medendorp, W. P. (2009). Spatial and effector processing in the human parietofrontal network for reaches and saccades. Journal of Neurophysiology, 101(6), 3053–3062. https://doi.org/10.1152/jn.91194.2008

Breveglieri, R., Borgomaneri, S., Filippini, M., Tessari, A., Galletti, C., Davare, M., & Fattori, P. (2022). Complementary contribution of the medial and lateral human parietal cortex to grasping: a repetitive TMS study. Cerebral Cortex. https://doi.org/10.1093/cercor/bhac404

Caminiti, R., Innocenti, G. M., & Battaglia-Mayer, A. (2015). Organization and evolution of parieto-frontal processing streams in macaque monkeys and humans. Neuroscience and Biobehavioral Reviews, 56, 73–96. https://doi.org/10.1016/j.neubiorev.2015.06.014

Castiello, U. (2005). The neuroscience of grasping. Nature Reviews. Neuroscience, 6(9), 726–736. https://doi.org/10.1038/nrn1744

Cavina-Pratesi, C., Connolly, J. D., Monaco, S., Figley, T. D., Milner, A. D., Schenk, T., & Culham, J. C. (2018). Human neuroimaging reveals the subcomponents of grasping, reaching and pointing actions. Cortex, 98, 128–148. https://doi.org/10.1016/j.cortex.2017.05.018

Cavina-Pratesi, C., Monaco, S., Fattori, P., Galletti, C., McAdam, T. D., Quinlan, D. J., Goodale, M. A., & Culham, J. C. (2010). Functional magnetic resonance imaging reveals the neural substrates of arm transport and grip formation in reach-to-grasp actions in humans. Journal of Neuroscience, 30(31), 10306–10323. https://doi.org/10.1523/JNEUROSCI.2023-10.2010

Comalli, D. M., Keen, R., Abraham, E. S., Foo, V. J., Lee, M. H., & Adolph, K. E. (2016). The development of tool use: Planning for end-state comfort. Developmental Psychology, 52(11), 878–1892. https://doi.org/10.1037/dev0000207

Culham, J. C., Cavina-Pratesi, C., & Singhal, A. (2006). The role of parietal cortex in visuomotor control: What have we learned from neuroimaging? Neuropsychologia, 44(13), 2668–2684. https://doi.org/10.1016/j.neuropsychologia.2005.11.003

Culham, J. C., Danckert, S. L., DeSouza, J. F. X., Gati, J. S., Menon, R. S., & Goodale, M. A. (2003). Visually guided grasping produces fMRI activation in dorsal but not ventral stream brain areas. Experimental Brain Research, 153(2), 180–189. https://doi.org/10.1007/s00221-003-1591-5

Culham, & Kanwisher. (2001). Neuroimaging of cognitive functions in human parietal cortex. Current Opinion in Neurobiology.

Di Bono, M. G., Begliomini, C., Castiello, U., & Zorzi, M. (2015). Probing the reaching-grasping network in humans through multivoxel pattern decoding. Brain and Behavior, 5(11), 1–18. https://doi.org/10.1002/brb3.412

Dijkerman, H. C., McIntosh, R. D., Schindler, I., Nijboer, T. C. W., & Milner, A. D. (2009). Choosing between alternative wrist postures: Action planning needs perception. Neuropsychologia, 47(6), 1476–1482. https://doi.org/10.1016/j.neuropsychologia.2008.12.002

Errante, A., Ziccarelli, S., Mingolla, G., & Fogassi, L. (2021). Grasping and Manipulation: Neural Bases and Anatomical Circuitry in Humans. Neuroscience, 458, 203–212. https://doi.org/10.1016/j.neuroscience.2021.01.028

Etzel, J. A., Zacks, J. M., & Braver, T. S. (2013). Searchlight analysis: Promise, pitfalls, and potential. In NeuroImage (Vol. 78, pp. 261–269). https://doi.org/10.1016/j.neuroimage.2013.03.041

Fabbri, S., Stubbs, K. M., Cusack, R., & Culham, J. C. (2016). Disentangling Representations of Object and Grasp Properties in the Human Brain. The Journal of Neuroscience, 36(29), 7648–7662. https://doi.org/10.1523/JNEUROSCI.0313-16.2016

Fischl, B., Sereno, M. I., Tootell, R. B. H., & Dale, A. M. (1999). High-Resolution Intersubject Averaging and aCoordinate System for the Cortical Surface. Human Brain Mapping.

Fogassi, L., Ferrari, P. F., Gesierich, B., Rozzi, S., Chersi, F., & Rizzolotti, G. (2005). Parietal lobe: From action organization to intention understanding. Science, 308(5722), 662–667. https://doi.org/10.1126/science.1106138

Frost, M. A., & Goebel, R. (2012). Measuring structural–functional correspondence: Spatial variability of specialised brain regions after macro-anatomical alignment. NeuroImage, 59(2), 1369–1381. https://doi.org/10.1016/j.neuroimage.2011.08.035

Gallivan, J. P., Adam McLean, D., Valyear, K. F., & Culham, J. C. (2013). Decoding the neural mechanisms of human tool use. ELife, 2013(2), 1–29. https://doi.org/10.7554/eLife.00425

Gallivan, J. P., Cant, J. S., Goodale, M. A., & Flanagan, J. R. (2014). Representation of object weight in human ventral visual cortex. Current Biology, 24(16), 1866–1873. https://doi.org/10.1016/j.cub.2014.06.046

Gallivan, J. P., Cavina-Pratesi, C., & Culham, J. C. (2009). Is that within reach? fMRI reveals that the human superior parieto-occipital cortex encodes objects reachable by the hand. Journal of Neuroscience, 29(14), 4381–4391. https://doi.org/10.1523/JNEUROSCI.0377-09.2009

Gallivan, J. P., Chapman, C. S., Gale, D. J., Flanagan, J. R., & Culham, J. C. (2019). Selective Modulation of Early Visual Cortical Activity by Movement Intention. Cerebral Cortex, 1–17. https://doi.org/10.1093/cercor/bhy345

Gallivan, J. P., & Culham, J. C. (2015). Neural coding within human brain areas involved in actions. In Current Opinion in Neurobiology (Vol. 33, pp. 141–149). Elsevier Ltd. https://doi.org/10.1016/j.conb.2015.03.012

Gallivan, J. P., Johnsrude, I. S., & Randall Flanagan, J. (2016). Planning Ahead: Object-Directed Sequential Actions Decoded from Human Frontoparietal and Occipitotemporal Networks. Cerebral Cortex, 26(2), 708–730. https://doi.org/10.1093/cercor/bhu302

Gallivan, J. P., McLean, D. A., Valyear, K. F., Pettypiece, C. E., & Culham, J. C. (2011). Decoding action intentions from preparatory brain activity in human parieto-frontal networks. Journal of Neuroscience, 31(26), 9599–9610.

Gamberini, M., Passarelli, L., Fattori, P., & Galletti, C. (2020). Structural connectivity and functional properties of the macaque superior parietal lobule. Brain Structure and Function, 225(4), 1349– 1367. https://doi.org/10.1007/s00429-019-01976-9

Gerbella, M., Rozzi, S., & Rizzolatti, G. (2017). The extended object-grasping network. Experimental Brain Research, 235(10), 2903–2916. https://doi.org/10.1007/s00221-017-5007-3

Goebel, R., Esposito, F., & Formisano, E. (2006). Analysis of Functional Image Analysis Contest (FIAC) data with BrainVoyager QX: From single-subject to cortically aligned group General Linear Model analysis and self-organizing group Independent Component Analysis. Human Brain Mapping, 27(5), 392–401. https://doi.org/10.1002/hbm.20249

Goodale, M. A., & Milner, D. A. (1992). Separate visual pathways for perception and action. Trends in Neurosciences.

Greenfield, A., Jahnsen, H., Downing, P. E., Jiang, Y., Shuman, M., & Kanwisher, N. (1996). A Cortical Area Selective for Visual Processing of the Human Body. In Trends Pharmacol. Sci (Vol. 16). https://www.science.org

Grefkes, C., & Fink, G. R. (2005). The functional organization of the intraparietal sulcus in humans and monkeys. 3–17.

Greve, D. N., & Fischl, B. (2009). Accurate and Robust Brain Image Alignment using Boundary-based Registration. Neuroimage, 48(1), 63–72. https://doi.org/10.1016/j.neuroimage.2009.06.060.Accurate

Guo, L. L., Nestor, A., Nemrodov, D., Frost, A., & Niemeier, M. (2019). Multivariate analysis of electrophysiological signals reveals the temporal properties of visuomotor computations for precision grips. Journal of Neuroscience, 39(48), 9585–9597. https://doi.org/10.1523/JNEUROSCI.0914-19.2019

Gutteling, T. P., Petridou, N., Dumoulin, S. O., Harvey, B. M., Aarnoutse, E. J., Leon Kenemans, J., & Neggers, S. F. W. (2015). Action preparation shapes processing in early visual cortex. Journal of Neuroscience, 35(16), 6472–6480. https://doi.org/10.1523/JNEUROSCI.1358-14.2015

Haxby, J. V., Gobbini, M. I., Furey, M. L., Ishai, A., Schouten, J. L., & Pietrini, P. (2001). Distributed and overlapping representations of faces and objects in ventral temporal corten. Science, 293(September), 87–96. https://doi.org/10.4324/9780203496190

Hinrichs, H., Scholz, M., Tempelmann, C., Woldorff, M. G., Dale, A. M., & Heinze, H. J. (2000). Deconvolution of event-related fMRI responses in fast-rate experimental designs: Tracking amplitude variations. Journal of Cognitive Neuroscience, 12. https://doi.org/10.1162/089892900564082

Kamitani, Y., & Tong, F. (2005). Decoding the visual and subjective contents of the human brain. Nature Neuroscience, 8(5), 679–685. https://doi.org/10.1038/nn1444

Kontaris, I., Wiggett, A. J., & Downing, P. E. (2009). Dissociation of extrastriate body and biological-motion selective areas by manipulation of visual-motor congruency. Neuropsychologia, 47(14), 3118–3124. https://doi.org/10.1016/j.neuropsychologia.2009.07.012

Kriegeskorte, N. (2008). Representational similarity analysis – connecting the branches of systems neuroscience. Frontiers in Systems Neuroscience, 2(November), 1–28. https://doi.org/10.3389/neuro.06.004.2008

Kriegeskorte, N., & Goebel, R. (2001). An efficient algorithm for topologically correct segmentation of the cortical sheet in anatomical MR volumes. NeuroImage, 14(2), 329–346. https://doi.org/10.1006/nimg.2001.0831

Kriegeskorte, N., & Kievit, R. A. (2013). Representational geometry: Integrating cognition, computation, and the brain. Trends in Cognitive Sciences, 17(8), 401–412. https://doi.org/10.1016/j.tics.2013.06.007

Monaco, S., Cavina-Pratesi, C., Sedda, A., Fattori, P., Galletti, C., & Culhaml, J. C. (2011). Functional magnetic resonance adaptation reveals the involvement of the dorsomedial stream in hand orientation for grasping. Journal of Neurophysiology, 106(5), 2248–2263. https://doi.org/10.1152/jn.01069.2010

Monaco, S., Malfatti, G., Culham, J. C., Cattaneo, L., & Turella, L. (2020). Decoding motor imagery and action planning in the early visual cortex: Overlapping but distinct neural mechanisms. NeuroImage, 218(May), 116981. https://doi.org/10.1016/j.neuroimage.2020.116981

Mylius, V., Ayache, S. S., Ahdab, R., Farhat, W. H., Zouari, H. G., Belke, M., Brugières, P., Wehrmann, E., Krakow, K., Timmesfeld, N., Schmidt, S., Oertel, W. H., Knake, S., & Lefaucheur, J. P. (2013). Definition of DLPFC and M1 according to anatomical landmarks for navigated brain stimulation: inter-rater reliability, accuracy, and influence of gender and age. NeuroImage, 78, 224–232. https://doi.org/10.1016/j.neuroimage.2013.03.061

Passarelli, L., Gamberini, M., & Fattori, P. (2021). The superior parietal lobule of primates: A sensory-motor hub for interaction with the environment. In Journal of Integrative Neuroscience (Vol. 20, Issue 1, pp. 157–171). IMR Press Limited. https://doi.org/10.31083/J.JIN.2021.01.334

Pertzov, Y., Avidan, G., & Zohary, E. (2011). Multiple reference frames for saccadic planning in the human parietal cortex. Journal of Neuroscience, 31(3), 1059–1068. https://doi.org/10.1523/JNEUROSCI.3721-10.2011

Picard, N., & Strick, P. L. (2001). Imaging the premotor areas. Current Opinion in Neurobiology, 11(6), 663–672. https://doi.org/10.1016/S0959-4388(01)00266-5

Pilacinski, A., Wallscheid, M., & Lindner, A. (2018). Human posterior parietal and dorsal premotor cortex encode the visual properties of an upcoming action. PLoS ONE, 13(10), 1–20. https://doi.org/10.1371/journal.pone.0198051

Pitzalis, S., Fattori, P., & Galletti, C. (2015). The human cortical areas V6 and V6A. In Visual Neuroscience (Vol. 32). Cambridge University Press. https://doi.org/10.1017/S0952523815000048

Ramon, M., Vizioli, L., Liu-Shuang, J., & Rossion, B. (2015). Neural microgenesis of personally familiar face recognition. Proceedings of the National Academy of Sciences of the United States of America, 112(35), E4835–E4844. https://doi.org/10.1073/pnas.1414929112

Rice, N. J., Valyear, K. F., Goodale, M. A., Milner, A. D., & Culham, J. C. (2007). Orientation sensitivity to graspable objects: An fMRI adaptation study. NeuroImage, 36(SUPPL. 2). https://doi.org/10.1016/j.neuroimage.2007.03.032

Richter, M., Amunts, K., Mohlberg, H., Bludau, S., Eickhoff, S. B., Zilles, K., & Caspers, S. (2019). Cytoarchitectonic segregation of human posterior intraparietal and adjacent parieto-occipital sulcus and its relation to visuomotor and cognitive functions. Cerebral Cortex, 29(3), 1305–1327. https://doi.org/10.1093/cercor/bhy245

Rizzolatti, G., & Matelli, M. (2003). Two different streams form the dorsal visual system: Anatomy and functions. Experimental Brain Research, 153(2), 146–157. https://doi.org/10.1007/s00221-003-1588-0

Rosenbaum, D. Α., Marchak, F., Barnes, H. J., Vaughan, J., Slotta, J. D., & Jorgensen, M. J. (1990). Constraints for Action Selection: Overhand Versus Underhand Grips. Attention and Performance XIII: Motor Representation and Control, August 2014, 321–342. https://doi.org/10.4324/9780203772010-10

Rosenke, M., van Hoof, R., van den Hurk, J., Grill-Spector, K., & Goebel, R. (2021). A Probabilistic Functional Atlas of Human Occipito-Temporal Visual Cortex. Cerebral Cortex, 31(1), 603–619. https://doi.org/10.1093/cercor/bhaa246

Scheperjans, F., Eickhoff, S. B., Hömke, L., Mohlberg, H., Hermann, K., Amunts, K., & Zilles, K. (2008). Probabilistic maps, morphometry, and variability of cytoarchitectonic areas in the human superior parietal cortex. Cerebral Cortex, 18(9), 2141–2157. https://doi.org/10.1093/cercor/bhm241

Scheperjans, F., Hermann, K., Eickhoff, S. B., Amunts, K., Schleicher, A., & Zilles, K. (2008). Observer-independent cytoarchitectonic mapping of the human superior parietal cortex. Cerebral Cortex, 18(4), 846–867. https://doi.org/10.1093/cercor/bhm116

Shikata, E., Hamzei, F., Glauche, V., Knab, R., Dettmers, C., Weiller, C., Büchel, C., & Büchel, B. (2001). Surface Orientation Discrimination Activates Caudal and Anterior Intraparietal Sulcus in Humans: An Event-Related fMRI Study. www.jn.physiology.org

Singhal, A., Monaco, S., Kaufman, L. D., & Culham, J. C. (2013). Human fMRI Reveals That Delayed Action Re-Recruits Visual Perception. PLoS ONE, 8(9). https://doi.org/10.1371/journal.pone.0073629

Tomassini, V., Jbabdi, S., Klein, J. C., Behrens, T. E. J., Pozzilli, C., Matthews, P. M., Rushworth, M. F. S., & Johansen-Berg, H. (2007). Diffusion-weighted imaging tractography-based parcellation of the human lateral premotor cortex identifies dorsal and ventral subregions with anatomical and functional specializations. Journal of Neuroscience, 27(38), 10259–10269. https://doi.org/10.1523/JNEUROSCI.2144-07.2007

Valyear, K. F., Culham, J. C., Sharif, N., Westwood, D., & Goodale, M. A. (2006). A double dissociation between sensitivity to changes in object identity and object orientation in the ventral and dorsal visual streams: A human fMRI study. Neuropsychologia, 44(2), 218–228. https://doi.org/10.1016/j.neuropsychologia.2005.05.004

van den Heiligenberg, F. M. Z., Orlov, T., Macdonald, S. N., Duff, E. P., Henderson Slater, D., Beckmann, C., Johansen-Berg, H., Culham, J. C., & Makin, T. R. (2018). Artificial limb representation in amputees. Brain, March, 1–12. https://doi.org/10.1093/brain/awy054

Vizioli, L., Bratch, A., Lao, J., Ugurbil, K., Muckli, L., & Yacoub, E. (2018). Temporal multivariate pattern analysis (tMVPA): A single trial approach exploring the temporal dynamics of the BOLD signal. Journal of Neuroscience Methods, 308(June 2018), 74–87. https://doi.org/10.1016/j.jneumeth.2018.06.029

Wandell, B. A., Dumoulin, S. O., & Brewer, A. A. (2007). Visual field maps in human cortex. In Neuron (Vol. 56, Issue 2, pp. 366–383). https://doi.org/10.1016/j.neuron.2007.10.012

Wood, D. K., Chouinard, P. A., Major, A. J., & Goodale, M. A. (2017). Sensitivity to biomechanical limitations during postural decision-making depends on the integrity of posterior superior parietal cortex. Cortex, 97, 202–220. https://doi.org/10.1016/j.cortex.2016.07.005

Yousry, T. A., Schmid, U. D., Alkadhi, H., Schmidt, D., Peraud, A., Buettner, A., & Winkler, P. (1997). Localization of the motor hand area to a knob on the precentral gyrus A new landmark. Brain, 141–157.

Zimmermann, M., Mars, R. B., de Lange, F. P., Toni, I., & Verhagen, L. (2018). Is the extrastriate body area part of the dorsal visuomotor stream? Brain Structure and Function, 223(1), 31–46. https://doi.org/10.1007/s00429-017-1469-0

Zimmermann, M., Verhagen, L., de Lange, F. P., & Toni, I. (2016). The extrastriate body area computes desired goal states during action planning. ENeuro, 3(2), 1918–1921. https://doi.org/10.1523/ENEURO.0020-16.2016

